# A tyrosine-based trafficking signal in the simian immunodeficiency virus envelope cytoplasmic domain is strongly selected for in pathogenic SIV infection

**DOI:** 10.1101/2021.03.31.437834

**Authors:** Scott P. Lawrence, Samra E. Elser, Workineh Torben, Robert V. Blair, Bapi Pahar, Pyone P. Aye, Faith Schiro, Dawn Szeltner, Lara A. Doyle-Meyers, Beth Haggarty, Andrea P.O. Jordan, Josephine Romano, Xavier Alvarez, David H. O’Connor, Roger W. Wiseman, Christine M. Fennessey, Yuan Li, Michael Piatak, Jeffrey D. Lifson, Celia C. LaBranche, Andrew A. Lackner, Brandon F. Keele, Nicholas J. Maness, Mark Marsh, James A. Hoxie

**Affiliations:** Perelman School of Medicine, University of Pennsylvania, Philadelphia, Pennsylvania, USA; MRC Laboratory for Molecular Cell Biology, University College London, London, United Kingdom; Tulane National Primate Research Center, Covington, Louisiana, USA; Wisconsin National Primate Research Center, Madison, Wisconsin, USA; AIDS and Cancer Virus Program, Frederick National Laboratory for Cancer Research, Frederick, Maryland; Duke University Medical Center, Durham, North Carolina, USA; Louisiana State University at Alexandria; Southwest National Primate Research Center, San Antonio, Texas, USA

**Keywords:** HIV, SIV, Envelope glycoprotein, trafficking, endocytosis, basolateral sorting

## Abstract

The HIV/SIV envelope glycoprotein (Env) cytoplasmic domain contains a highly conserved Tyr-dependent trafficking signal that mediates both clathrin-dependent endocytosis and polarized sorting of Env. Despite extensive characterization, the role of these functions in viral infection and pathogenesis is unclear. An SIV molecular clone (SIVmac239) in which the Tyr-based signal is inactivated by deletion of Gly-720 and Tyr-721 (SIVmac239ΔGY) replicates to high levels acutely in pigtail macaques (PTM) but is rapidly controlled. We previously reported that rhesus macaques and PTM can progress to AIDS following SIVmac239ΔGY infection in association with novel amino acid changes in the Env cytoplasmic domain. These included an R722G flanking the ΔGY deletion and a nine nucleotide deletion that encodes amino acids 734-736 (ΔQTH) and overlaps with the *rev* and *tat* open reading frames. We show that molecular clones containing these mutations reconstitute signals for both endocytosis and polarized sorting. In one PTM, a novel genotype was selected, which generated a new signal for polarized sorting but not endocytosis. This mutation by itself was sufficient to maintain high viral loads for several months when introduced into naïve PTMs. These findings reveal, for the first time, strong selection pressure for Env endocytosis and, in particular, for polarized sorting during pathogenic SIV infection *in vivo*.

## INTRODUCTION

The central themes that underlie pathogenesis of human immunodeficiency virus type-1 (HIV-1) in humans, and simian immunodeficiency virus (SIV) in Asian macaques, include failure of the host to control viral replication, chronic immune activation, and progressive loss of CD4+ T cells across multiple anatomic sites (1–3). Although the disease AIDS can take years to develop, there are critical early events that dictate the outcome of infection. CD4+/CCR5+ T cells in gut-associated lymphoid tissue (GALT) are massively depleted within a month of initial infection leading to compromised epithelial barrier function, microbial translocation and systemic immune activation (1–5). There are also pathological innate sensors engaged within hours of SIV infection that dysregulate antiviral interferon responses and initiate proinflammatory programs that are sustained and may compromise subsequent adaptive immune responses (6–10). Nevertheless, immune control of viral replication can occur, although the mechanisms and determinants for this outcome are poorly understood (11–13).

The envelope glycoprotein (Env) of HIV-1 and SIV is expressed on the surface of infected cells and on virions as a trimer of gp120/gp41 subunits in which gp41 anchors Env to cellular and viral membranes. Cellular infection is initiated when viral gp120 binds to CD4 and a coreceptor (CCR5 or CXCR4), leading to gp41 mediated membrane fusion (14). A conserved feature of gp41 is a long cytoplasmic domain of approximately 160 amino acids, which contains a number of motifs that engage cellular trafficking machineries and signaling pathways (15, 16). We have shown that deletion of Gly and Tyr (a.a. 720 and 721) from a highly conserved GYxxØ-type trafficking motif (x= any a.a.; Ø= an a.a. with a bulky hydrophobic side chain) (17) in the Env of SIVmac239 (GYRPV), creates a virus (termed SIVmac239ΔGY) that in pigtail macaques (PTM) leads to an extraordinary phenotype of acute plasma viral RNA peaks similar to parental SIVmac239, but is subsequently controlled (<15-50 RNA copies/ml) following peak viremia with the onset of host cellular immune responses (18). In contrast to SIVmac239 infection, in SIVmac239ΔGY infection, CD4+ T cells are not depleted in the blood or gut, infection of GALT is only transient, there is no microbial translocation or chronic immune activation, and animals remain healthy as elite controllers for months to years (18). Host control of SIVmac239ΔGY is independent of neutralizing antibodies and associated with strong, polyfunctional antiviral CD4+ and CD8+ T cell responses with a role for CD8+ T cells (CTL or NK cells) shown by anti-CD8 cell depletion (18). Interestingly, in rhesus macaques (RM), control of SIVmac239ΔGY infection is incomplete, and animals progress to disease with a detectable viral load. The fact that PTM can suppress viral replication completely is paradoxical, given that PTM typically progress to AIDS more rapidly than do RM following SIVmac239 infection (19).

For HIV-1 and SIV Envs the GYxxØ motif, together with a less conserved C-terminal di-leucine motif, influence the expression and distribution of Env on infected cells by engaging clathrin-based trafficking pathways (17, 20–22). Through a direct interaction with AP1 and AP2 clathrin adaptor complexes, these motifs maintain low levels of Env on the surface of infected cells that are likely to contribute to the well recognized paucity of Env on virions (22–24). In addition, the HIV-1 Env GYxxØ motif (GYSPL), in the context of a full viral genome, has been shown to direct polarized sorting of Env to the basal and lateral plasma membrane in MDCK cells (25–28), and the SIV and HIV-2 GYxxØ motifs (GYRPV) shown to direct polarized sorting of Env in Vero cells (27). HIV-1 Env has been suggested to influence the polarized sorting of the viral Gag protein in T cells (25), although Gag multimerization independent of Env has also been implicated in polarized viral budding in T cells (29–31). Regardless, the *in vivo* roles for Env endocytosis and polarized sorting are unclear, as are the mechanisms through which deletion of GY from the GYRPV trafficking signal in SIVmac239 has such a profound impact on persistence and viral pathogenesis.

Here we evaluated the effects of the ΔGY deletion on viral assembly and replication *in vitro* and assessed the effects of previously reported mutations acquired in macaques (RM and PTM) that were infected with SIVmac239ΔGY but progressed to disease (32, 33). These mutations included an R722G change flanking the ΔGY deletion and loss of 3 downstream amino acids, QTH (a.a. 734-736 [ΔQTH]). We show that ΔGY decreased Env content within cells and on virions, which is restored by R722G. Remarkably, ΔQTH generated new Tyr-dependent signals for both Env endocytosis and polarized sorting. When introduced into the SIVmac239ΔGY Env, these mutations could partially restore pathogenesis in PTM, and in some animals were associated with persistent viremia and progression to AIDS. In an additional animal inoculated with a SIVmac239ΔGY virus containing R722G that progressed to AIDS, 3 point mutations appeared in the cytoplasmic domain of Env that generated a new signal for polarized sorting, but not endocytosis, despite also creating a stop codon in the second exon of *tat*. When introduced into the SIVmac239ΔGY+R722G virus and inoculated into PTM, all 3 point mutations were retained throughout infection and were sufficient to confer sustained intermediate to high levels of viremia for several months. Together these results indicate that a reduction in Env on virions and loss of cellular trafficking functions caused by the ΔGY deletion could be restored *in vivo* by acquisition of novel compensatory mutations and that reacquisition of these functions correlated with a gain of pathogencity. In particular, our findings reveal strong selection pressures to maintain polarized trafficking of Env *in vivo* and demonstrate that a loss of this function can lead to potent host immune control.

## RESULTS

### Mutations acquired during pathogenic ΔGY infection

We previously reported that 4 of 4 RM (32) and 2 of 21 PTM (18) infected with SIVmac239ΔGY progressed to AIDS. Single genome amplification (SGA) and sequencing revealed that the ΔGY deletion was maintained in all viral amplicons from plasma from these animals, in addition it identified novel mutations in the Env cytoplasmic domain. These included either a R722G substitution flanking the ΔGY deletion, which restored a Gly at a.a. position 720, or an S727P substitution (Figure 1). In 2 RM, acquisition of R722G was followed by extraordinary deletions of 9 nucleotides (nt 8803-8811 in animal DT18 and nt 8804-8812 in animal DD84) that removed a.a. 734-736 (QTH) generating a YFQI and YFQL a.a. sequence, respectively. These mutations are unique among SIVmac-related sequences listed in the Los Alamos HIV Sequence Database and remarkable in that (i) they generated a YxxØ sequence reminiscent of the conserved GYRPV motif disrupted by the ΔGY deletion, and (ii) the 9 nucleotide deletions occured within the overlapping reading frames for the second exons of *tat* and *rev* (Supplemental Figure 1). The S727P substitution had been seen in an earlier study of SIVmac239ΔGY infection in RM (34) and was shown to increase infection in gut CD4+ T cells during acute infection (33). Given that either R722G (with or without ΔQTH) or S727P was seen in all 6 animals infected with SIVmac239ΔGY that progressed to disease (18, 32), we evaluated the impact of these changes on Env expression and trafficking *in vitro* and on pathogenesis *in vivo*.

**Figure 1.**
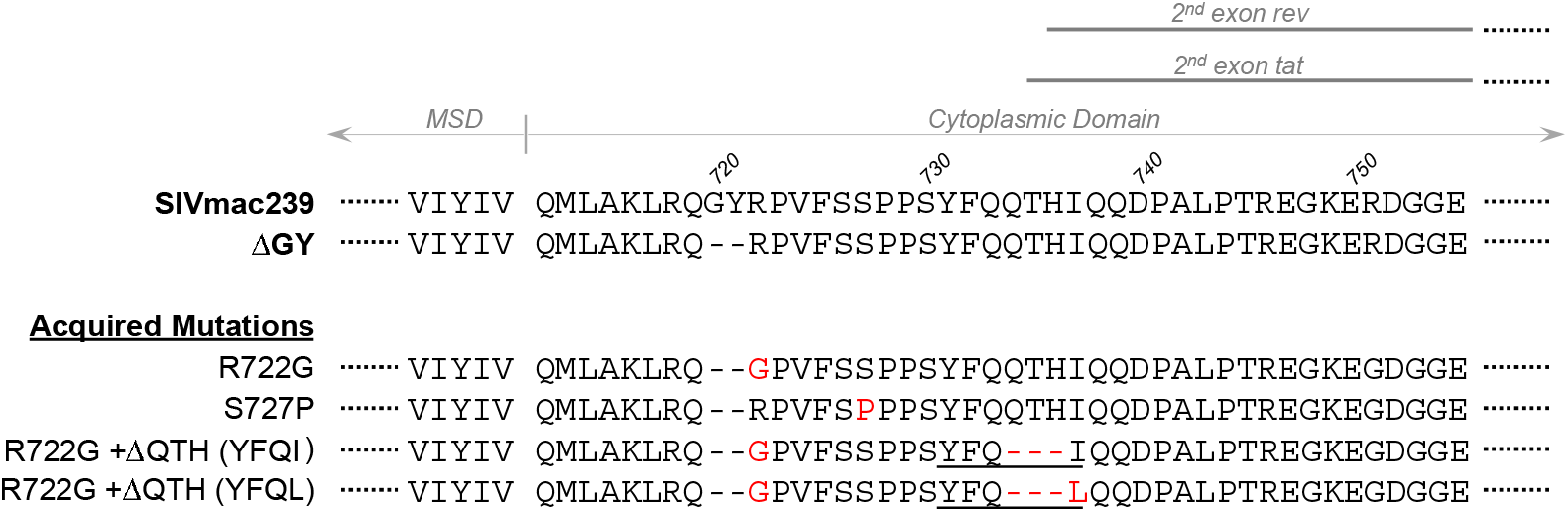
Amino acid changes acquired in Env during pathogenic SIVmac239ΔGY infection. Amino acid (a.a.) sequences of SIVmac239 and ΔGY Envs indicating the membrane spanning domain (MSD), the predicted start of the cytoplasmic domain, and approximate start sites for the second exons of *tat* and *rev* in overlapping reading frames. Below (in red) are a.a. changes in this region previously reported in rhesus (32) and pigtail macaques (18) that failed to control ΔGY and progressed to disease. YxxØ motifs created by ΔQTH mutations are underlined (see Supplemental Figure 1 for nt sequences).

### Effects of ΔGY and ΔGY-associated changes on Env expression

To characterize the effects of the ΔGY deletion and the mutations acquired *in vivo*, we first assessed Env expression on infected cells and virions by quantitiative western blotting (Figure 2). To avoid any variation in particle infectivity due to different SIV Envs, we used VSV-G pseudotyped SIV with genomes encoding either SIVmac239 or ΔGY Envs ± ΔGY-associated mutations. Cell lines lacking CD4 were used to avoid cytopathic effects during Env expression.

**Figure 2.**
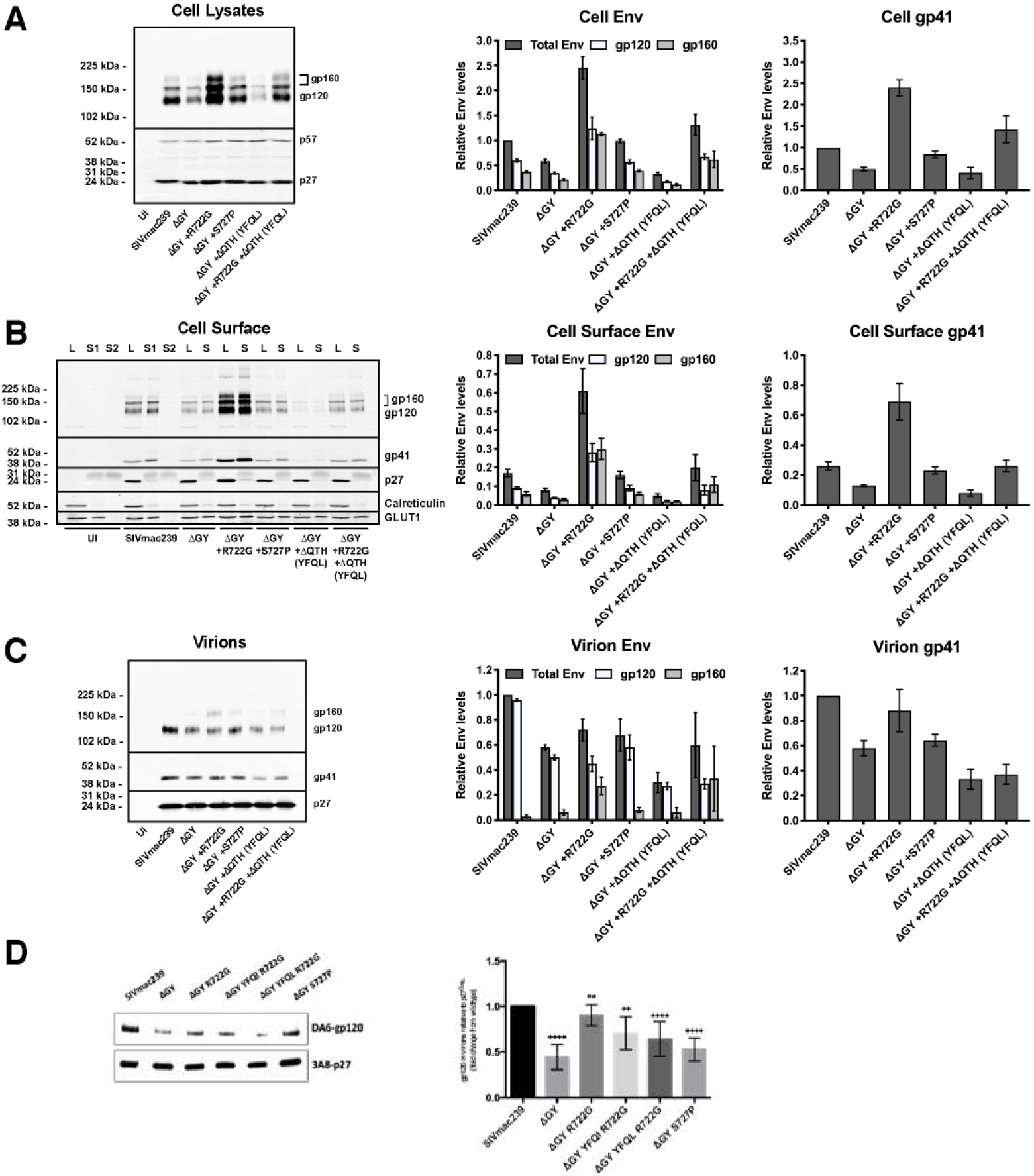
The ΔGY deletion and *in vivo* mutations modulate Env content on cells and virions. Expression of total cell, cell surface and virion-associated Env was determined by western blotting using LLC-MK2 cells infected with viruses encoding the indicated Envs. **(A)** Total cell-associated Env is shown for cell lysates. **(B)** Cell surface Env isolated by biotinylation and pull-down is shown (S = Surface). Cell lysates (L = lysate) correspond to 20% of the pull-down input. A second pull-down (S2) was performed to show that all biotinylated proteins were captured in the first round. **(C)** Env associated with virions released into supernatant from cells shown in Panel A. **(D)** Env associated with virions produced from infected rhesus macaque PBMCs. Left panels show representative western blots; right panels show quantitation of the western blots. Env levels are normalized to cell p57/p27 and relative to SIVmac239 set at 1. The intensity of the processed (gp120) or unprocessed Env (gp160) is shown as the fraction of the total Env signal for each lane and relative to SIVmac239 set at 1. Graphs display the mean ± SEM from n ≥ 3 independent experiments.

#### Effects of the ΔGY mutation on Env content in cells and on virions

Relative to Gag p57/p27, ΔGY Env content in total cell lysates and on the cell surface of rhesus LLC-MK2 cells was reduced approximately 40% compared to SIVmac239 Env (Figure 2A and B). The decrease in Env was similar for both mature Env (gp120/gp41) and gp160, indicating that Env cleavage was unaltered. Relative to Gag p27, ΔGY Env content on virions was reduced approximately 40% compared to SIVmac239 Env (Figure 2C). Similar results were seen for both gp120 and gp41, indicating that Env shedding could not explain this difference. Levels of uncleaved gp160 on virions were negligible for SIVmac239, although a slight increase from 3% to 11% of the total Env (P=0.0193) was seen on ΔGY virions. Thus, the reduced level of ΔGY Env on virions corresponded to a general decrease in cellular ΔGY Env levels.

#### Effects of ΔGY-associated mutations acquired *in vivo* on Env content in cells and on virions

Mutations encoding the amino acid changes shown in Figure 1 were introduced into SIVmac239ΔGY and Env levels were determined in cells and on virions as above. Strikingly, the reduction in Env content (gp120 and gp160) caused by the ΔGY deletion was rescued by R722G with expression exceeding the levels of total cell-associated and cell surface SIVmac239 Env (Figure 2A and B). R722G also increased ΔGY Env on virions and in cells when it contained the ΔQTH (YFQL) mutation. On virions, this increase was predominantly due to gp160, suggesting less efficient cleavage of Env prior to incorporation into virions. Indeed, the addition of R722G to a ΔGY Env background resulted in a small reduction of Env processing (18.7% ± 4.5, P=0.02) when compared to SIVmac239-infected cells. S727P had a negligible effect on virion Env levels but, when introduced into ΔGY Env, restored Env content in infected cells to levels of SIVmac239 (Figure 2A, B and C). In contrast, a ΔQTH mutation (producing YFQL) reduced total cell-associated and cell surface Env content, as well as virion Env, when introduced into a ΔGY background with or without R722G (Figure 2A, B and C). Similar, results were seen for all mutants when experiments were performed using HEK293T cells and the human T lymphoid cell lines BC7 and CEMx174 (data not shown).

#### Effects of ΔGY-associated mutations on envelope content on virions produced in primary T cells

To determine the effects of ΔGY and acquired Env changes on virions produced in primary macaque lymphocytes, RM and PTM PBMCs, were activated using Conconavalin A and IL2 and infected with viruses containing SIVmac239 or ΔGY Envs, or ΔGY Envs containing R722G, R722G + ΔQTH changes (producing either YFQI or YFQL), or S727P substitution. Virions were harvested after 4 days and analyzed for Env content by western blotting. As shown in Figure 2D for RM PBMCs, relative to virion-associated p27-Gag, ΔGY Env was reduced approximately 50% compared to SIVmac239, similar to that seen in virus produced in LLC-MK2 cells (Figure 2B). As in LLC-MK2 cells, R722G restored levels of Env containing the ΔGY deletion to near SIVmac239 levels, while S727P had a lesser effect.

Thus, while the ΔGY deletion resulted in a decrease in Env content in ΔGY-infected cells, on the cell surface, and on virions, this defect was largely corrected by the R722G substitution. In contrast, ΔQTH reduced Env content on cells and virions relative to Envs containing ΔGY alone or with R722G.

### Alterations in the cellular distribution of Env caused by the ΔGY deletion and acquired mutations

The ΔGY deletion is expected to ablate a trafficking signal for clathrin-dependent endocytosis with the potential to alter the cellular distribution of Env (17, 21, 22, 35). To assess the effects of ΔGY and the mutations acquired *in vivo*, we used previously described chimeric reporter constructs that contain the CD4 ecto- and membrane spaning domains (MSD) fused to the SIV Env cytoplasmic domain (CD) (Figure 3A) (22, 35). Because there are additional endocytosis signals in SIV and HIV Envs distal to the GYxxØ motif (22, 35, 36), these constructs contained only the membrane proximal 30 a.a. of the SIVmac Env CD that lack additional endocytic trafficking information (22). HeLa cell lines were generated that stabily expressed these CD4-based constructs containing SIVmac239 or ΔGY CDs with or without the changes described in Figure 1.

**Figure 3.**
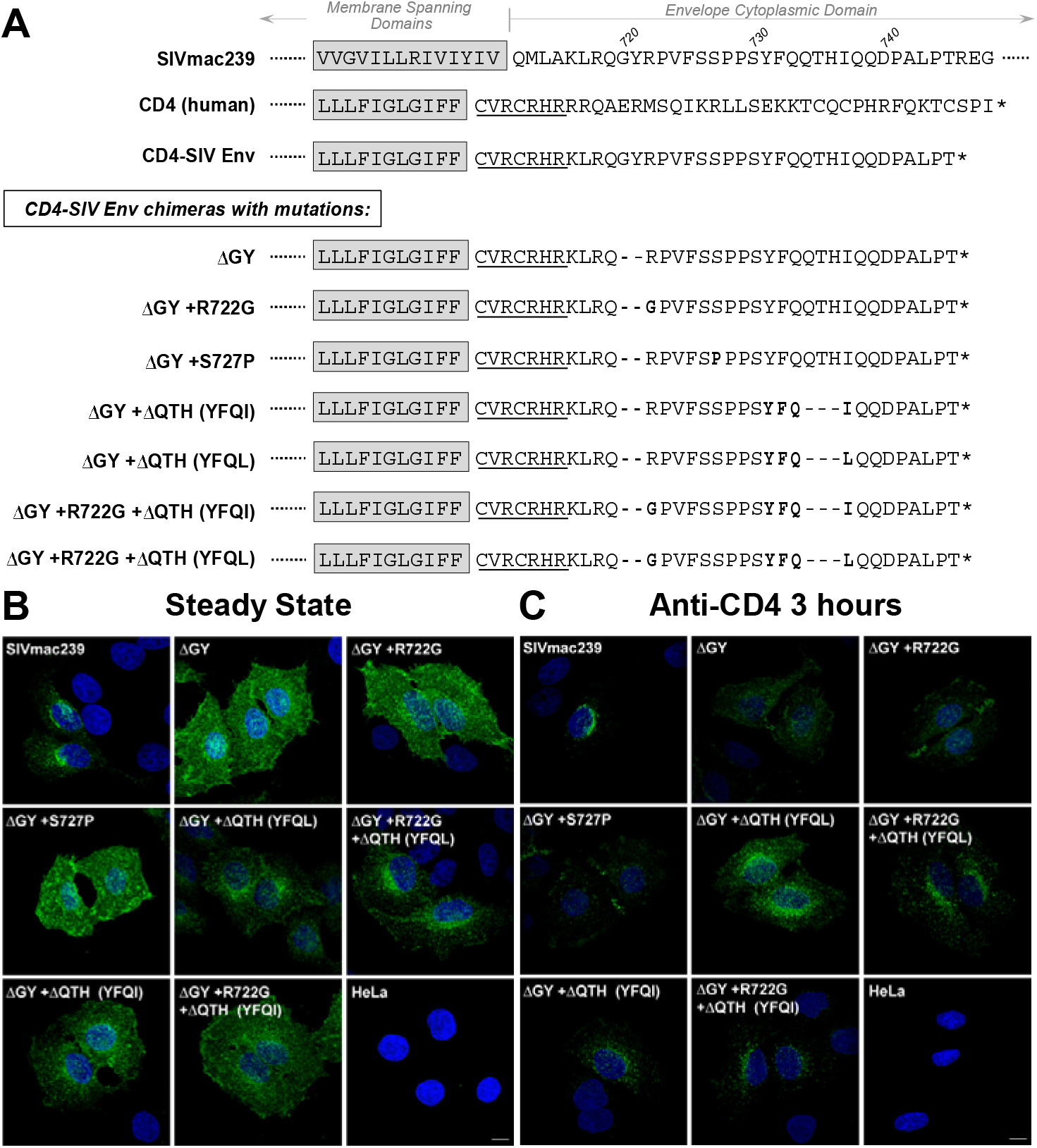
Alterations in cellular trafficking induced by ΔGY and *in vivo* acquired mutations. **(A)** Partial sequences for SIVmac239 Env and human CD4 are shown with the membrane spanning domains (shaded) and cytoplasmic domains (CD; partial for SIVmac239 Env and full length for CD4). A CD4-SIV Env CD chimeric construct is shown containing the CD4 ecto- and membrane spanning domains with 7 a.a. from the membrane proximal cytoplasmic domain of CD4 (underlined) fused to a.a. 716-745 from the SIVmac239 Env CD (20, 22). CD4-SIV Env CD chimeras are shown containing ΔGY and indicated changes (**bold**). **(B)** Steady state cellular distribution of the indicated CD4-SIV Env CD chimeras stably expressed in HeLa cells. **(C)** Cellular distribution of CD4-SIV Env CD chimeras following incubation of cells with anti-CD4 antibody (Q4120) at 37°C for 3 hrs prior to fixation. Confocal Z stacks were deconvolved and displayed as maximum projections. Size bar = 10 μm.

The steady state distribution of the constructs was evaluated on fixed and permeabilized cells using an anti-CD4 antibody (22). The chimera containing the SIVmac239 CD was detected predominantly in a perinuclear pattern consistent with an intracellular distribution, while the construct containing the ΔGY CD was diffusely distributed on the cell surface (Figure 3B). When the ΔGY CD contained ΔQTH, the clustered perinuclear pattern was restored, while ΔGY CD chimeras containing the R722G or S727P substitutions remained diffusely distributed on the cell surface. To determine if any constructs trafficked to the cell surface and were then internalized, cells were incubated with anti-CD4 antibody for 3 hours at 37°C prior to fixation and permeabilization. Cells expressing the SIVmac239 Env CD chimera exhibited prominent punctate intracellular staining showing that this protein had been exposed on the cell surface and then endocytosed, while the ΔGY CD construct remained predominantly on the cell surface (Figure 3C). Strikingly, when the ΔGY CD chimera contained the ΔQTH deletions, perinuclear punctate patterns were again seen, while addition of R722G or S727P to the ΔGY CD chimera had no effect. Collectively, these findings indicated that endocytic trafficking functions of the SIV CD that had been ablated by the ΔGY mutation were restored by ΔQTH, but not by the R722G or S727P substitutions.

### ΔQTH deletions create novel Tyr-dependent endocytosis signals

We quantified the effects of the ΔQTH mutation on the endocytic properties of CD4-SIV Env CD constructs shown in Figure 3 by measuring the uptake of an anti-CD4 antibody over time using a modification of a previously described protocol (22). Cells were incubated with antibody at 4°C, washed and warmed to 37°C; the decrease of cell-surface antibody was then measured over time (Figure 4). Endocytosis of the SIVmac239 CD construct followed the same two phase pattern previously described (22); during the rapid phase (0-5 min) of endocytosis, where recycling is negligible, the SIVmac239 Env CD construct was endocytosed at ~12% per minute, whereas endocytosis of the ΔGY Env was reduced to 1.8 ± 0.6% per minute, a rate consistent with bulk membrane turnover (Figure 4A). While addition of R722G or S727P to the ΔGY CD had only a negligible effect on endocytic rates, the ΔQTH deletion that generated the YFQI sequence, partially restored the endocytic rate to 5.7 ± 0.4% per minute, while the ΔQTH deletion that generated YFQL restored the endocytic rate to SIVmac239 levels (12.8 ± 0.6%) (Figure 4A).

**Figure 4.**
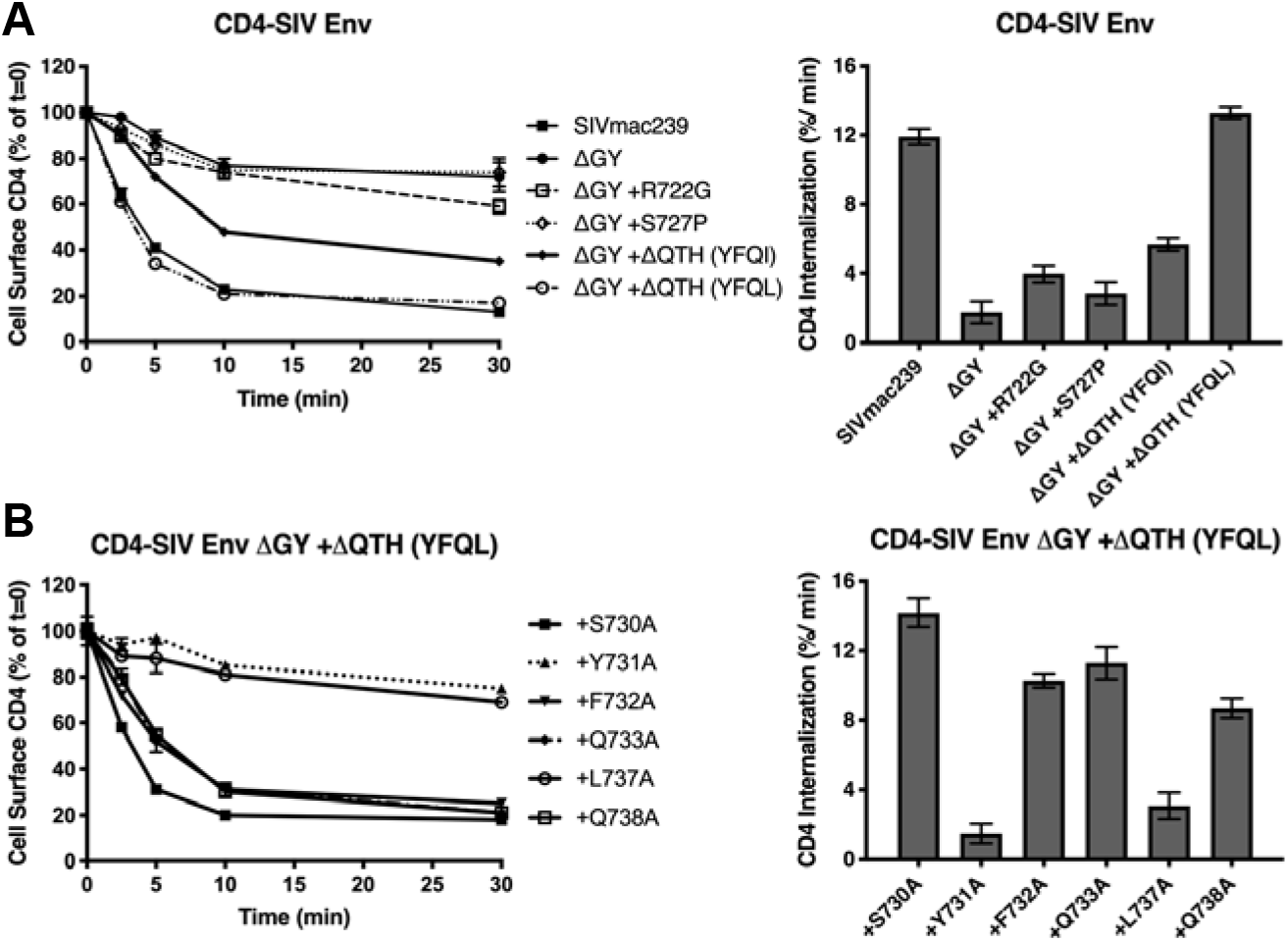
ΔGY and *in vivo* acquired mutations modulate endocytic rates. **(A)** Endocytosis of CD4-SIV Env CD chimeras shown in Figure 3A in HeLa cells. **(B)** Endocytosis of CD4-SIV Env CD constructs containing ΔGY + ΔQTH (YFQL) with the indicated alanine substitutions (SIVmac239 a.a. numbering). Left panels show cell surface CD4 for the indicted times as a % of 0 mins (=100%); right panels show the rate of endocytosis over the first 5 minutes after warm up. Graphs display the mean ± S.E.M. from ≥ 3 independent experiments.

To determine if the YFQI and YFQL sequences created endocytosis signals that conformed to a conventional YxxØ motif, alanine substitutions were introduced into CD4-SIV Env CD constructs that contained ΔGY and the ΔQTH deletion that generated YFQL, and endocytosis rates were determined on stably-expressing HeLa cells (Figure 4B). Substitution of Y731A (position 0 of YFQL) or L737A (position Y+3) reduced endocytosis to ΔGY levels, whereas substitutions at the Y+1, +2 and +4 positions had only minor effects, consistent with a classical YxxØ endocytic signal (37). The Y731A substitution also altered the cellular distribution of a CD4-SIV Env CD construct from predominantly intracellular to the cell surface (Supplemental Figure 2A and B). To further determine if the YxxØ motifs (YFQI and YFQL) exhibited features typical of a classical endocytosis motif, AP2 μ2 subunits were depleted with siRNA and the endocytic rates of CD4-SIV Env CD chimeras measured. Depletion of μ2 has been shown to destabilize the AP2 complex and ablate endocytic function (38). Efficient μ2 depletion was achieved (69 ± 12%), and destabilization of the AP2 complex demonstrated with a 53 ± 13% reduction of the α-adaptin subunit (Supplemental Figure 3A and B). Notably, AP2 knock down reduced the internalization rates of SIV Env CD chimeras containing the parental GYRPV or the YFQI or YFQL sequences associated with the ΔQTH mutations (Supplemental Figure 3C).

Thus, both ΔQTH deletions that occurred in the Env CDs of SIVmac239ΔGY-infected macaques that progressed to AIDS regenerated endocytosis signals that fully restored the endocytic function of the parental SIVmac239 sequence (YRPV) ablated by the ΔGY mutation, and were confirmed to be authentic, AP2-dependent YxxØ endocytosis signals.

### ΔQTH mutations also create novel basolateral sorting signals

For both HIV and SIV Envs, the Tyr in the membrane proximal GYxxØ motif has been shown to mediate basolateral sorting of Env expressed in polarized epithelial cells (25, 27, 28, 39). To determine if the YFQI and YFQL motifs generated by ΔQTH deletions also reconstituted basolateral sorting. CD4-SIV Env CD constructs with or without these changes (Figure 3A) were stably expressed in MDCKII cells, which polarize to form apical and basolateral surfaces when cultured as monolayers (40). The panel of constructs used included CD4-SIV Env CD chimeras from SIVmac239 with truncated (short tail) or full length (long tail) CDs (Figure 5A and B, respectively). Surface expression of CD4-SIV Env constructs containing either SIVmac239 short or full-length CDs was low but localized to the basolateral membrane as visualized by microscopy and quantified by determining the basolateral/apical distribution ratio (Figure 5A and B). Introduction of Y721I or ΔGY resulted in complete loss of this polarization (Figure 5A and B), consistent with previous findings that basolateral sorting of Env is dependent on Y721 (25). In contrast to the presence of multiple endocytic signals in the Env CD (22), these results indicated that polarized sorting of the CD4-SIV Env CD chimera was determined solely by the Tyr-containing motif. In agreement with these observations, a basolateral distribution was also seen for native, full length SIVmac239 Env, which was also ablated by the ΔGY mutation (Figure 5C). R722G or S727P substitutions in the CD4-SIV Env short tail construct containing ΔGY did not restore polarized sorting (Figure 5A). However, when the ΔQTH deletions were introduced, sorting to the basolateral surface was fully restored with an increase in the basolateral/apical ratio comparable to the SIVmac239 CD construct. Similarly, a Y731A mutation in the YFQL sequence, completely ablated this distribution (Figure 5A). Thus, the new YxxØ motifs, YFQI and YFQL, created by the ΔQTH deletions, completely restored Tyr-dependent polarized sorting CD4-SIV Env CD constructs that contained the ΔGY deletion.

**Figure 5.**
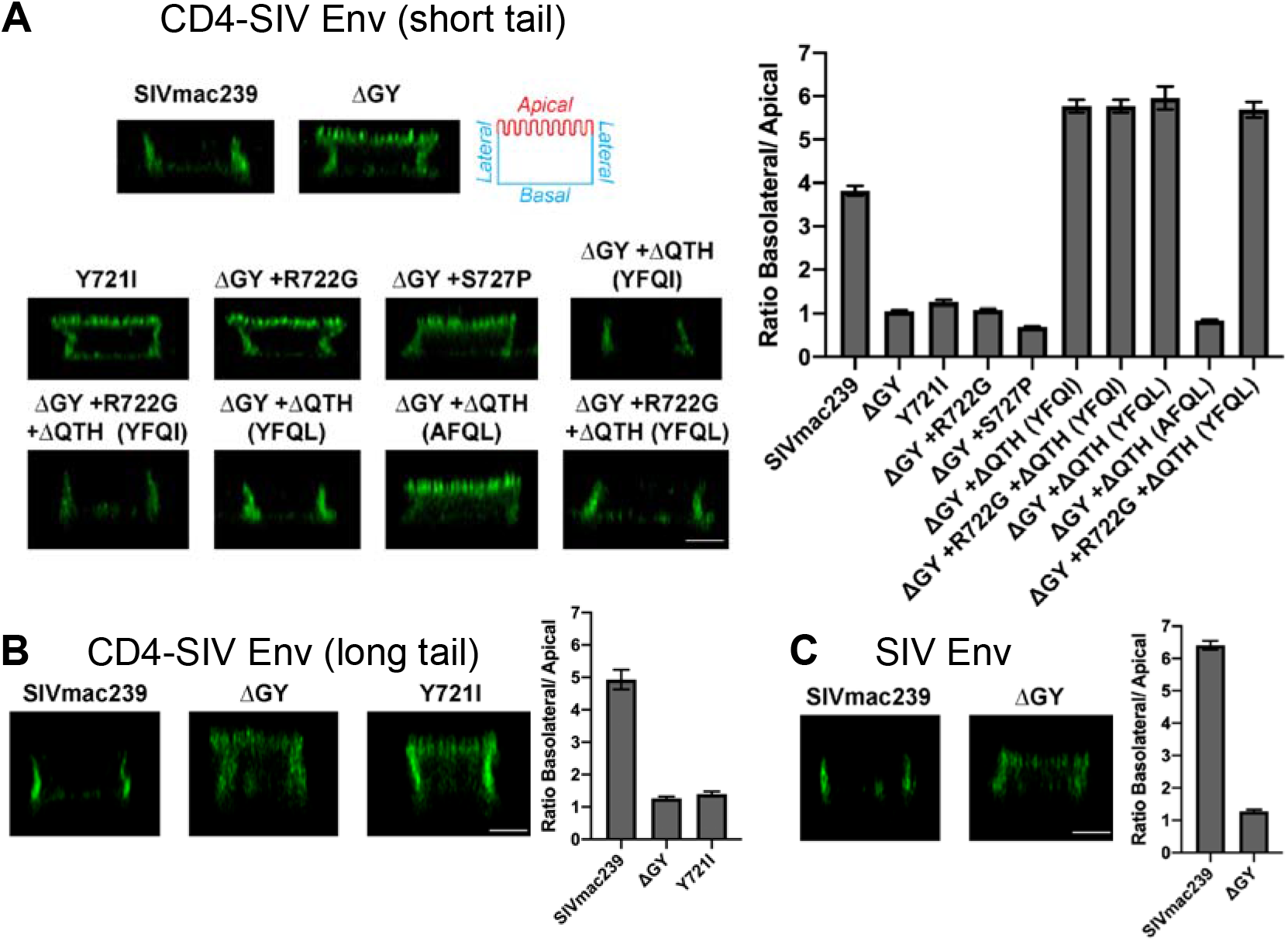
ΔGY and *in vivo* acquired mutations modulate Env sorting in polarized epithelial cells. **(A)** Cell surface distribution of CD4-SIV Env short tail chimeras containing the indicated Env CDs from Figure 3 on polarized MDCKII cells. Apical, basal, and lateral surfaces are indicated. **(B)** Cell surface distribution of CD4-SIV Env chimeras containing a full-length SIVmac239 CD (long tail). **(C)** Cell surface distribution of native, full length SIVmac239 Env with or without the ΔGY mutation. Left panels show an orthogonal deconvolved projection of a representative cell; right panels show quantitation of the images. Data shown is the average of between 58-223 cells per condition imaged from **(A)** n≥3 [except ΔGY + ΔQTH (AFQL) which is n=1] **(B)** n≥2 and **(C)** n=2 independent experiments. Scale bar = 5 μm.

### Evaluating the effects of the R722G substitution and ΔQTH deletions *in vivo*

We sought to determine the impact of the R722G substitution and ΔQTH deletions on SIVmac239ΔGY pathogenesis *in vivo*. SIVmac239ΔGY viruses containing R772G, with or without the ΔQTH deletions (generating either YFQI or YFQL), were replication competent *in vitro* in PTM PBMCs, although SIVmac239ΔGY viruses containing ΔQTH deletions lacking R722G replicated poorly (Supplemental Figure 4). Two viruses were selected for *in vivo* studies, one containing ΔGY with both R722G and the ΔQTH deletion generating YFQL (designated SIVmac239ΔGY+R722G +ΔQTH) and the other containing ΔGY with R722G alone (designated SIVmac239ΔGY+R722G). We selected PTM for this evaluation given the usually potent viral control and absence of disease in this species following SIVmac239ΔGY infection (18). Two groups of 3 PTM were inoculated i.v. with 300 TCID_50_ of virus, and animals followed for plasma viremia, CD4+ T cells in blood and gut, and the stability of mutations over time by single genome sequencing of plasma viral RNA (32, 33).

#### Infection by SIVmac239ΔGY containing both R722G and ΔQTH resulted in persisting viremia

The 3 animals inoculated with SIVmac239ΔGY containing both R722G and the ΔQTH deletion (KV74, KV52, and KV76) exhibited acute peak viral loads in plasma ranging from 4.6 - 6.0×10^6^ copies/ml, (Figure 6A). Although the levels of viral RNA varied during chronic infection, all animals maintained detectable viremia for 65 weeks with KV74 having high and stable levels (0.4 to 1.7×10^5^ copies/ml), KV52 showing a gradual decline, and KV76 showing marked fluctuations (levels ranging from 0.3 - 2.2×10^4^ RNA copies/ml). Gut CD4+ T cells initially declined for all animals during the first 2-4 weeks of infection to levels that were lower than for historical ΔGY-infected PTMs, but then recovered with KV74 remaining at ~50% of preinfection levels, and KV76 and KV52 showing a gradual return to baseline (Supplemental Figure 5). KV74 also showed a reduction in platelets, a recognized indicator of AIDS in PTM (41, 42), between 16 and 40 weeks post-infection in combination with low levels of CD4 T cells comparable to SIVmac239 (Supplemental Figure 5 and 6), although platelets in this animal subsequently recovered to baseline (Supplemental Figure 6).

**Figure 6.**
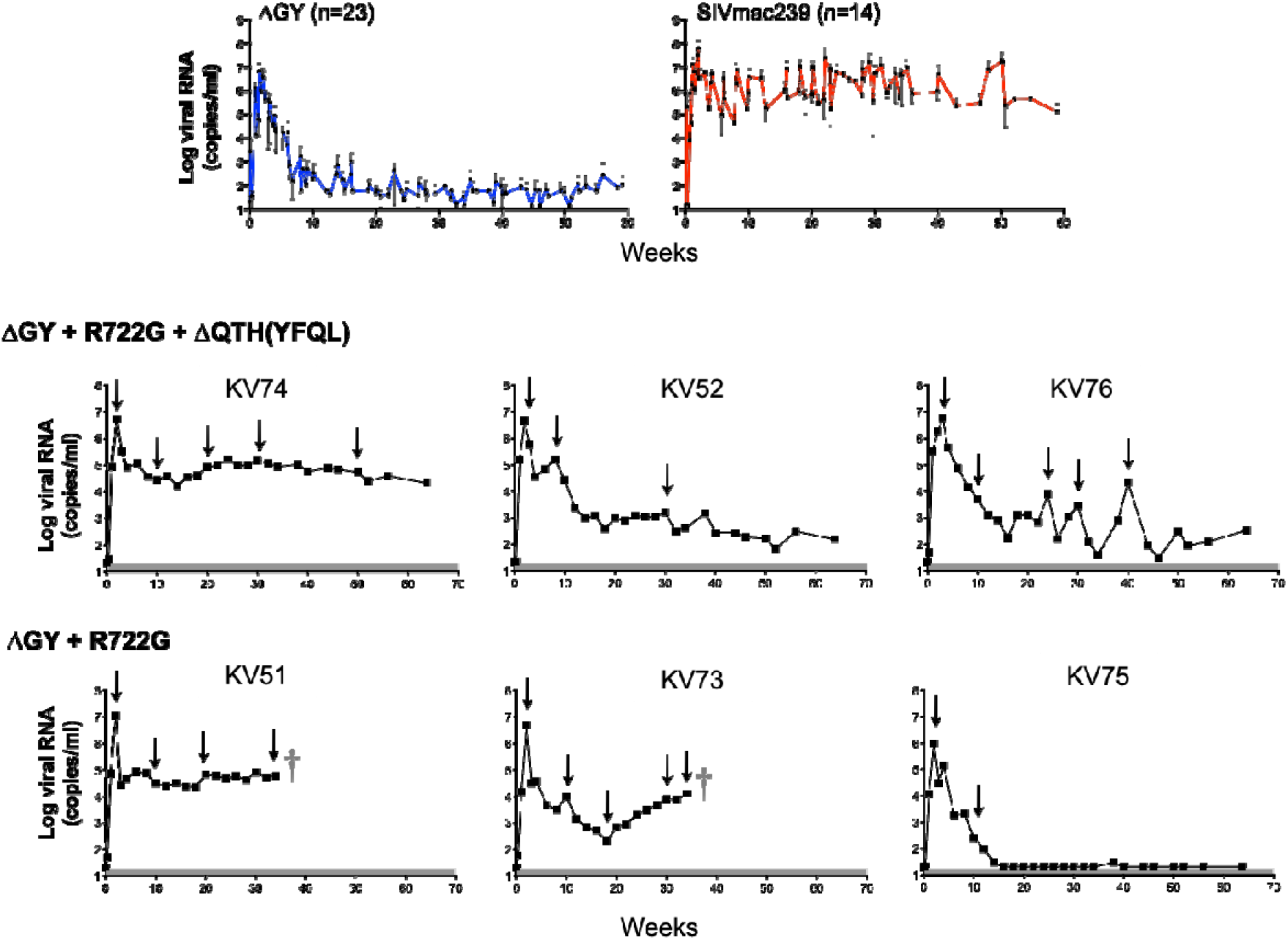
Plasma viral loads for pigtail macaques infected with SIVmac239ΔGY containing mutations acquired *in vivo*. Top panels show plasma viral loads from historical PTM controlling SIVmac239ΔGY (n=23; left panel) and SIVmac239-infected controls (n=14; right panel) (18) (and unpublished results). Lower panels show plasma viral loads for individual animals inoculated with ΔGY containing R722G and the ΔQTH deletion that generated YFQL (animals KV52, KV74 and KV76) and ΔGY containing R722G alone (animals KV51, KV73 and KV75). † indicates death due to an AIDS-related complication. Arrows denote time points at which plasma was obtained for SGA analysis of viral sequences. Shaded areas indicate approximate limits of sensitivity for the assay, which varied from 15-50 copies/ml for animals shown.

SGA and sequencing of viral RNA from plasma was performed at multiple time points (Figure 6). The sequences for the Env CD are shown for all amplicons (Supplemental Figure 7A-C) with a summary of changes shown for Env a.a. 719-752 encompasing ΔGY, R722G, and ΔQTH (Table 1) and Env a.a. 860-880 encompassing the distal CD (Supplemental Table 1). SGA sequencing showed persistence of the R722G and ΔQTH deletion in all amplicons and at all time points with the exception of 1 of 25 amplicons in KV74 at week 50 in which the ΔQTH was lost (Table 1 and Supplemental Figure 7A-C). Persistence of the ΔQTH deletion was remarkable given that the nucleic acid sequence generating ΔQTH creates deletions within the *tat* and *rev* second exons (Supplemental Figure 1). An analysis of *tat* and *rev* mRNA transcripts in PBMCs from all 3 animals assessed 14 days post infection confirmed these deletions were present *in vivo* and that alternative splice acceptors were being used (Supplemental Figure 8).

**Table 1.**
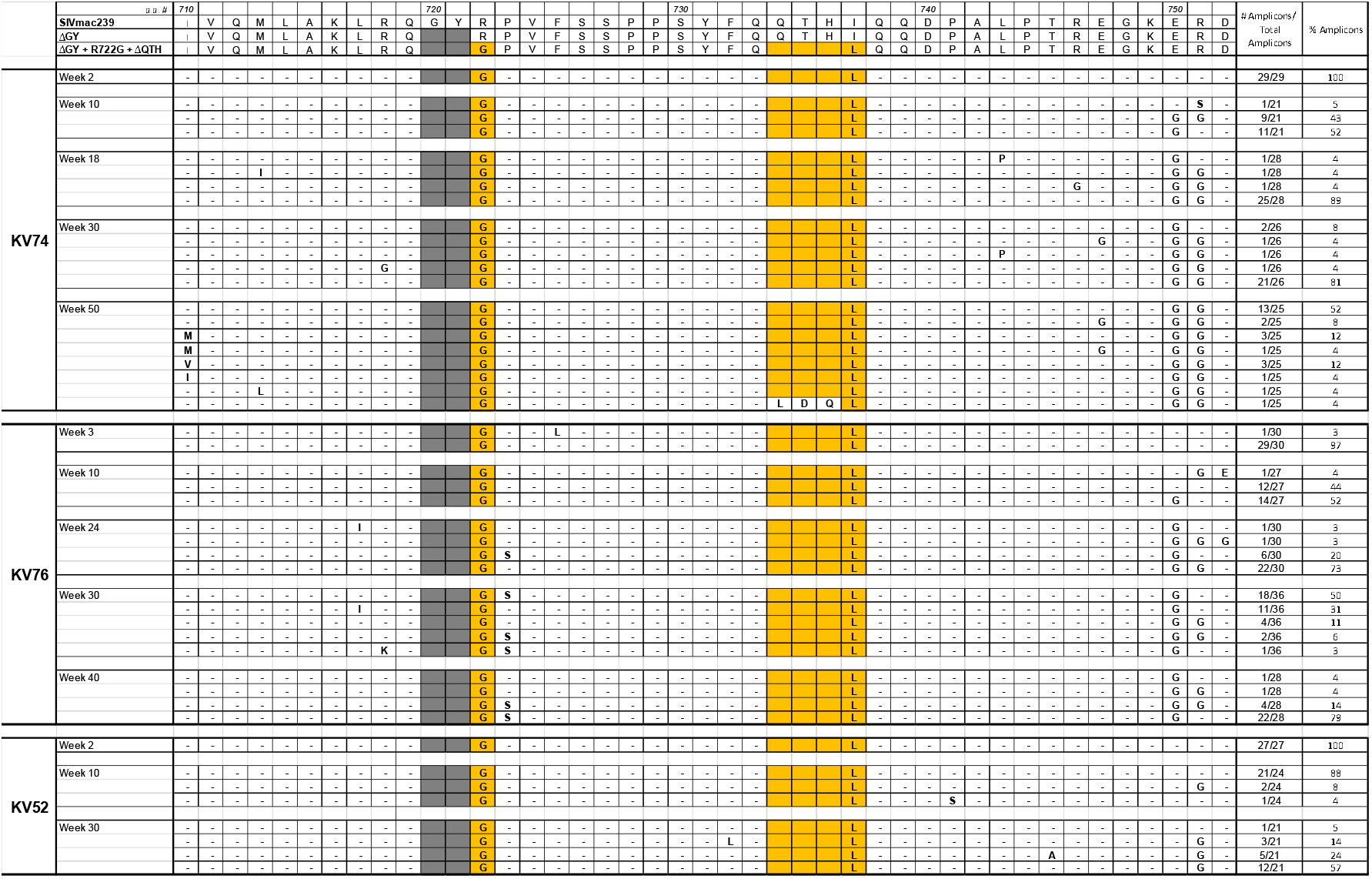
Viral evolution in macaques inoculated with SIVmac239ΔGY+R722G+ΔQTH. Summary of single genome sequencing of plasma virus from 3 pigtail macaques inoculated with SIVmac239ΔGY+R722G+ΔQTH. Results are shown for mutations within Env a.a. 710-752. The numbers and % of amplicons bearing the indicated mutations are shown. Dashes indicate identity with SIVmac239ΔGY+R722G+ΔQTH; shaded area indicates a ΔGY deletion. Sequences in the top panel show Env of SIVmac239, ΔGY and ΔGY+R722G+ΔQTH from parent viruses. Colored boxes show mutations introduced into the SIVmac239ΔGY background and their conservation over time in each animal. The ΔQTH deletion that was introduced also generated an I737L substitution as shown. The Env cytoplasmic domain sequences of individual amplicons is given in Supplemental Figure 7.

A well described fitness mutation for parental SIVmac239 (R751G) (43), appeared in all animals by week 10, with or without an adjacent E750G substitution (Supplemental Figure 7A-C). Although less commonly observed *in vivo*, E750G has been reported previously in SIVmac239-infected cynomolgus macaques (44). In animal KV76, a P723S substitution, not seen in other animals, appeared at week 24 and was nearly fixed by week 34 (Table 1 and Supplemental Figure 7B). Additional sporadic changes were seen in these animals including V815A and V837A, and in the C-terminus G873E or R, L874I, and L879S, although none of these were fixed (Supplemental Table 1 and Supplemental Figure 7A-C). Thus, the addition of R722G and ΔQTH to SIVmac239ΔGY resulted in a high level of sustained viremia in one animal and variable but persistent levels of viremia in two others for over 1 year. These findings are in marked contrast to SIVmac239ΔGY infection where viral control typically occurs within 8-10 weeks (Figure 6) (18). Importantly, complete retention of R722G and ΔQTH indicated that there was strong selection pressure to maintain these changes and their functions *in vivo*.

#### Infection with SIVmac239ΔGY+R722G leads to disease progression in association with new changes in the Env cytoplasmic domain

The 3 animals inoculated with SIVmac239ΔGY+R722G (KV51, KV73 and KV75) exhibited acute viral peaks at week 2 of 1.1×10^7^, 5.2×10^6^, and 9.8×10^5^ RNA copies/ml, respectively (Figure 6). In KV75, the viral load decreased to 100 copies/ml by week 12 and thereafter became undetectable. In this animal, gut CD4+ T cells decreased from 50 to 18% of T cells at week 2, but increased to preinfection levels with the decline in viremia (Supplemental Figure 5). In contrast, KV51 poorly controlled virus with viremia persisting between 2.3-8.5×10^4^ copies/ml, while in KV73, viremia decreased to a low of 210 copies/ml at week 18, but thereafter increased to 1.3×10^4^ copies/ml by week 35. Gut CD4+ T cells for both of these animals decreased to <5% at the time of peak viremia, with KV51 remaining at 25% of preinfection levels and KV75 returning to its preinfection level by week 28 (Supplemental Figure 5). Remarkably, KV51 and KV73 both developed severe thrombocytopenia (Supplemental Figure 6) and died at weeks 36 and 37, respectively, with massive pulmonary artery thrombi, a complication frequently seen in PTM during pathogenic SIV infection (18, 45–48).

SGA and and sequencing of plasma virus for KV75, at weeks 2 and 10 showed that the ΔGY deletion and R722G substitution were maintained (Table 2 and Supplemental Figure 7D) and at week 10 both R751G and E750G appeared. Only one additional change in Env appeared, an A836D substitution in 19 of 20 amplicons (Supplemental Figure 7D, Supplemental Table 2), which is contained within a CTL epitope targeted by both RM (49) and PTM (50). In marked contrast, KV51 and KV73 exhibited striking new changes that emerged during progression to disease. In KV51, the ΔGY and R722G changes were maintained, but at week 24 an in-frame 9 nt deletion (nt 8803-8011) appeared in 26 of 29 amplicons (Table 2 and Supplemental Figures 7E and F) that generated a ΔQTH deletion and YFQI sequence (Figure 1 and Supplemental Figures 1 and 8). In 7 amplicons, a YFQL sequence resulting from an I737L substitution, was also found. By week 34, all 26 amplicons contained ΔQTH, 24 with YFQL and 2 with YFQI, indicating a selection advantage for YFQL (Table 2). Distal to the R751G substitution with or without an adjacent E750G, no other changes were seen in >10% amplicons with the exception of an L874I substitution near the Env C-terminus at week 34 (Supplemental Table 2 and Supplemental Figure 7E).

**Table 2.**
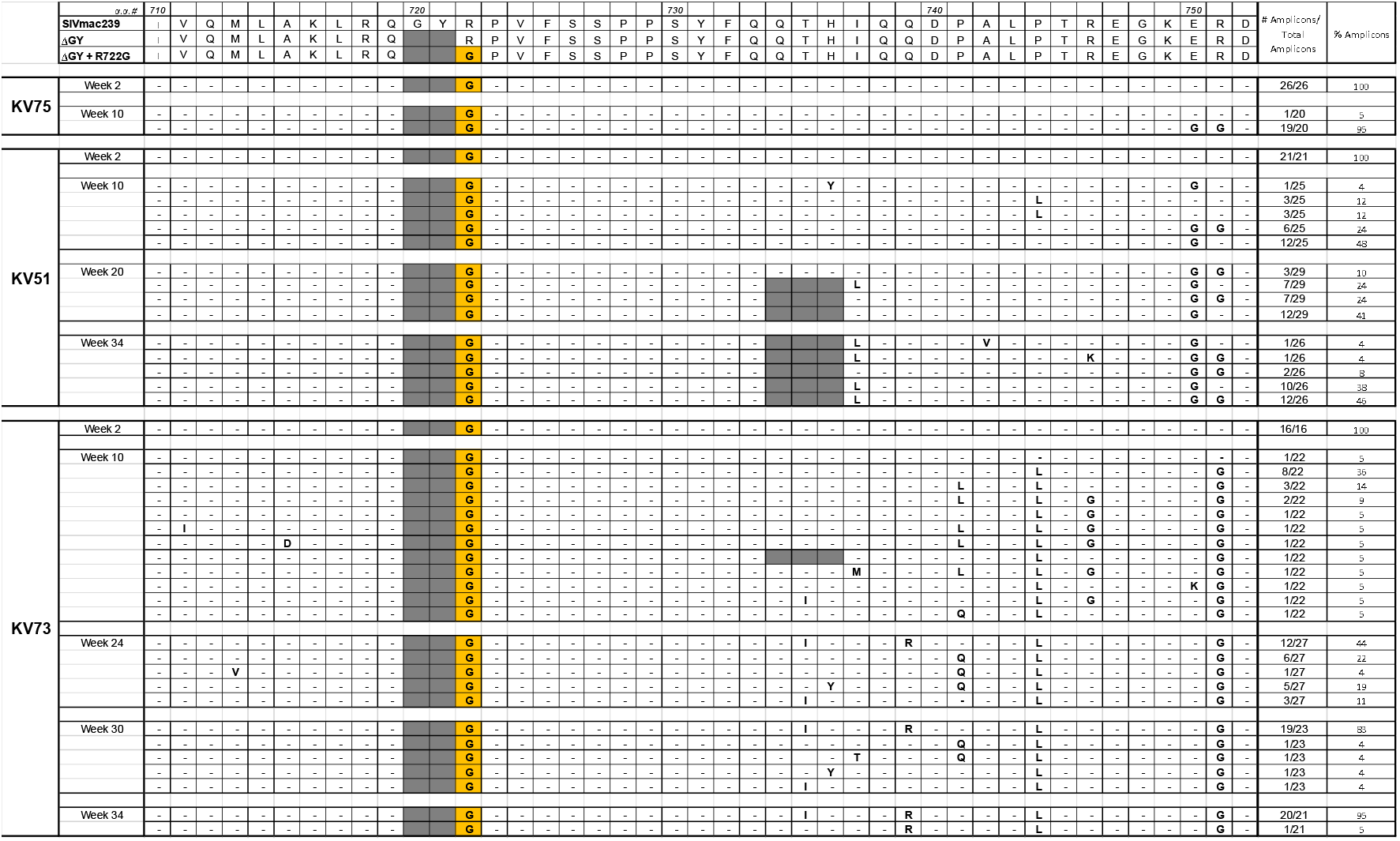
Viral evolution in macaques inoculated with SIVmac239ΔGY containing the R722G substitution. Summary of single genome amplification and sequencing of plasma virus performed at the indicated time points for 3 pigtail macaques inoculated i.v. with SIVmac239ΔGY+R722G. Results are shown for changes within a.a. 710-752 as in Table 1. Sequences in the top panel show Env of SIVmac239, SIVmac239ΔGY and SIVmac239ΔGY+R722G. Complete a.a. sequences in the cytoplasmic domain for individual amplicons are shown in Supplemental Figure 7.

In KV73, the Env ΔGY and R722G substitution were also conserved throughout infection with R751G again appearing by week 10 (Table 2 and Supplemental Figure 7F). Interestingly, a ΔQTH deletion appeared in 1 of 22 amplicons at week 10 but was lost at all subsequent time points. However, starting at week 10, a.a. substitutions were observed between positions 735 and 744 of Env (Table 2) including T735I, H736Y, Q739R, P741Q or L, and P744L. The proportions of these changes varied, but evolved by week 34 to a consensus of T735I, Q739R, and P744L in 20 of 21 amplicons (Table 2), henceforth termed the “IRL” mutation set. Based on publicly available sequences, T735I and Q739R have been observed individually during SIVmac239 infection but never together, while P744L has not been reported. As shown, (Supplemental Figure 9) nt mutations that created the T735I and H736Y in Env, along with a G to A mutation that is silent in Env, generated 3 coding mutations in the *tat* second exon. Remarkably, P744L created a stop codon in the second exon of *tat* causing a deletion of the last 22 a.a., which was verified by RT-PCR and sequencing of viral mRNA transcripts when a virus containing these mutations was grown *in vitro* in PTM PBMCs. V837A and G873E also appeared at week 10 and evolved to become fixed mutations (Supplemental Table 2 and Supplemental Figure 7F).

Thus, in the two animals infected with SIVmac239ΔGY+R722G that progressed to AIDS, each evolved new changes in the Env CD: in KV51, a ΔQTH deletion generating YFQL; and in KV73, the IRL set. The previous appearance of ΔQTH in RM and now in PTM again suggested strong selection pressure to restore trafficking signals for Env endocytosis and polarized sorting. However, the IRL set did not correspond to any recognized cellular trafficking signal.

### The IRL set confers a novel signal for Env polarized sorting but not endocytosis

To determine if the IRL set acquired in animal KV73 during progression to AIDS influenced Env trafficking, these mutations, individually and in combination, were introduced into CD4-SIV Env CD chimeras (Figure 7A). Constructs containing ΔGY and R722G with IRL or single amino acid changes within IRL (T735I, Q739R, or P744L) showed no increase in endocytosis rates compared to ΔGY and displayed minimal uptake of anti-CD4 antibody into intracellular compartments (Figure 7B and Supplemental Figure 2). In contrast, ΔGY +R722G Env CD chimeras containing all 3 IRL substitutions showed potent basolateral sorting in MDCKII cells, equivalent to that of SIVmac239 Env CD (Figure 7C). Individual T735I and P744 substitutions partially reconstituted basolateral sorting, although all 3 were required for maximal effect. These results indicate that *in vivo* there is strong selection pressure to generate polarized trafficking function. Moreover, the finding that the IRL set, in contrast to ΔQTH, restored polarized sorting but not endocytic function suggests that, at least for the membrane proximal GYxxØ motif, polarized trafficking of Env is the more important of these functions *in vivo*.

**Figure 7.**
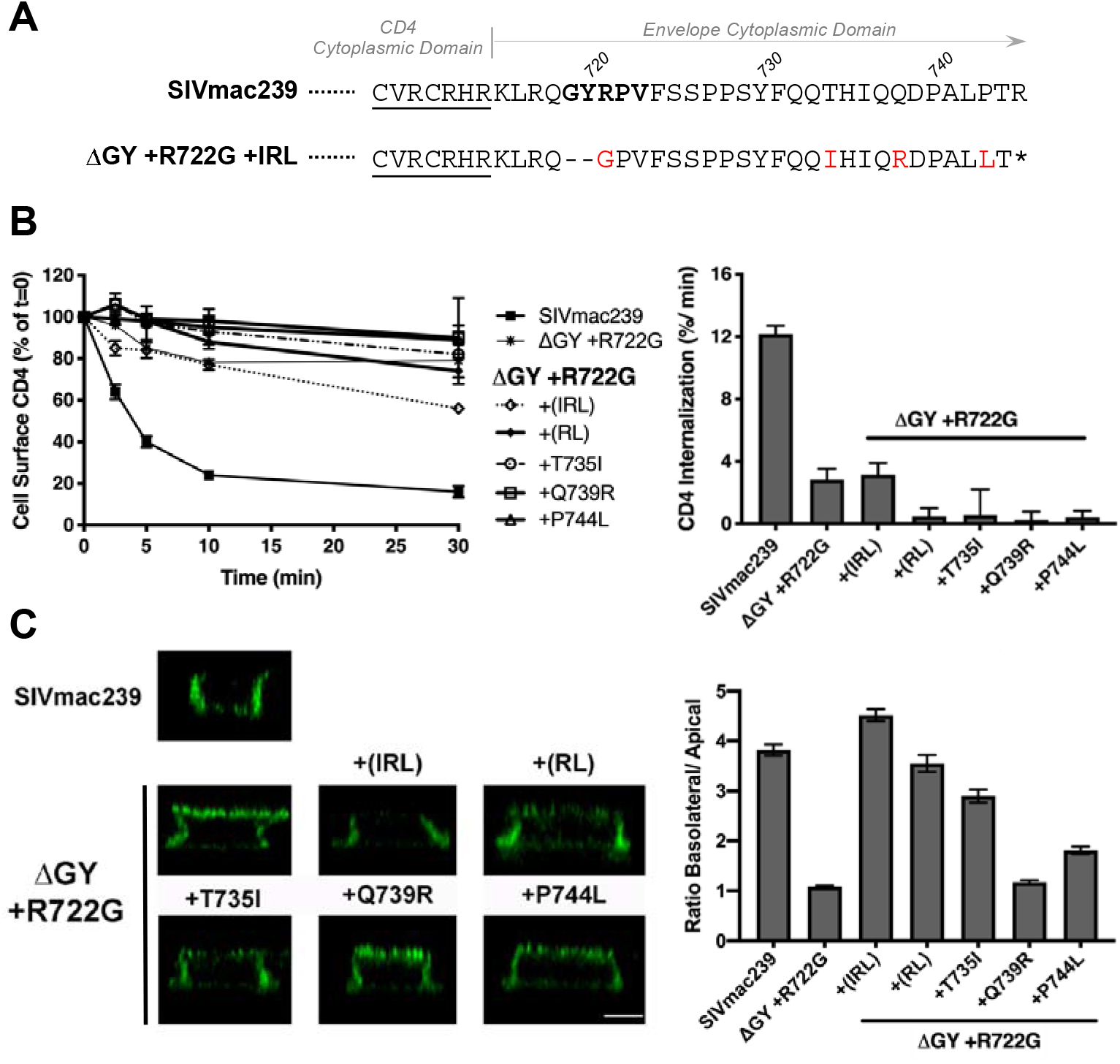
The IRL substitutions restore polarized sorting of Env but not endocytosis. **(A)** The C-terminal sequences of CD4-SIV Env CD chimeric constructs (Figure 3A) are shown with sequences from the CD4 CD underlined. Constructs containing the WT SIVmac239 CD or this CD containing ΔGY, R722G, and IRL substitutions are shown (red). **(B)** Endocytosis of CD4-SIV Env short tail chimeras in HeLa cells. Left panel shows the amount of cell surface CD4 at the indicted times as a % of 0 mins (=100%); right panel shows the rate of CD4 endocytosis over the first 5 mins after warm-up. Graphs display the mean ± SEM from 3 independent experiments. **(C)** Cell surface CD4-SIV Env CD chimeras on polarized MDCKII cells. Left panels show orthogonal deconvolved immunofluorescence projections of representative cells; right panel shows quantitation of the ratio of basolateral to apical signal from the left panels. The data are averages of between 59-190 cells per condition from n≥2 independent experiments. Scale bar = 5 μm.

### The polarized sorting function of the IRL set is conserved *in vivo* and confers persistent elevated levels of viremia to SIVmac239ΔGY

We determined if the IRL changes were sufficient to restore pathogenicity to a virus containing the ΔGY mutation. IRL substitutions (T735I, Q739R and P744L) were introduced into the SIVmac239ΔGY+R722G virus, given the absolute conservation of R722G *in vivo*, and inoculated i.v. into 4 PTM. These animals (NH85, NH86, NH87 and NH88) exhibited intermediate to high acute viral peaks of 2.3×10^7^, 3.4×10^7^, 2.7×10^5^, and 1.7×10^6^, respectively, and maintained elevated levels of viremia for up to 30 weeks of 1.3×10^3^, 2.5×10^3^, 3.8×10^4^, and 5.1×10^5^ copies/ml (Figure 8). Thereafter, NH85 and NH86 decreased to <100 copies/ml, while NH87 and NH88 increased to terminal values at week 40 of 9.2×10^4^ and 2.3×10^5^ copies/ml, respectively. Gut CD4+ T cells transiently decreased in 3 animals, although recovery to preinfection levels occurred by week 20 (Supplemental Figure 5). The persistent viremia in these animals was in marked contrast to historical PTM inoculated with SIVmac239ΔGY, which exhibited viral set points typically <15-50 copies/ml by 10-20 weeks of infection (18) (see also Figure 6).

**Figure 8.**
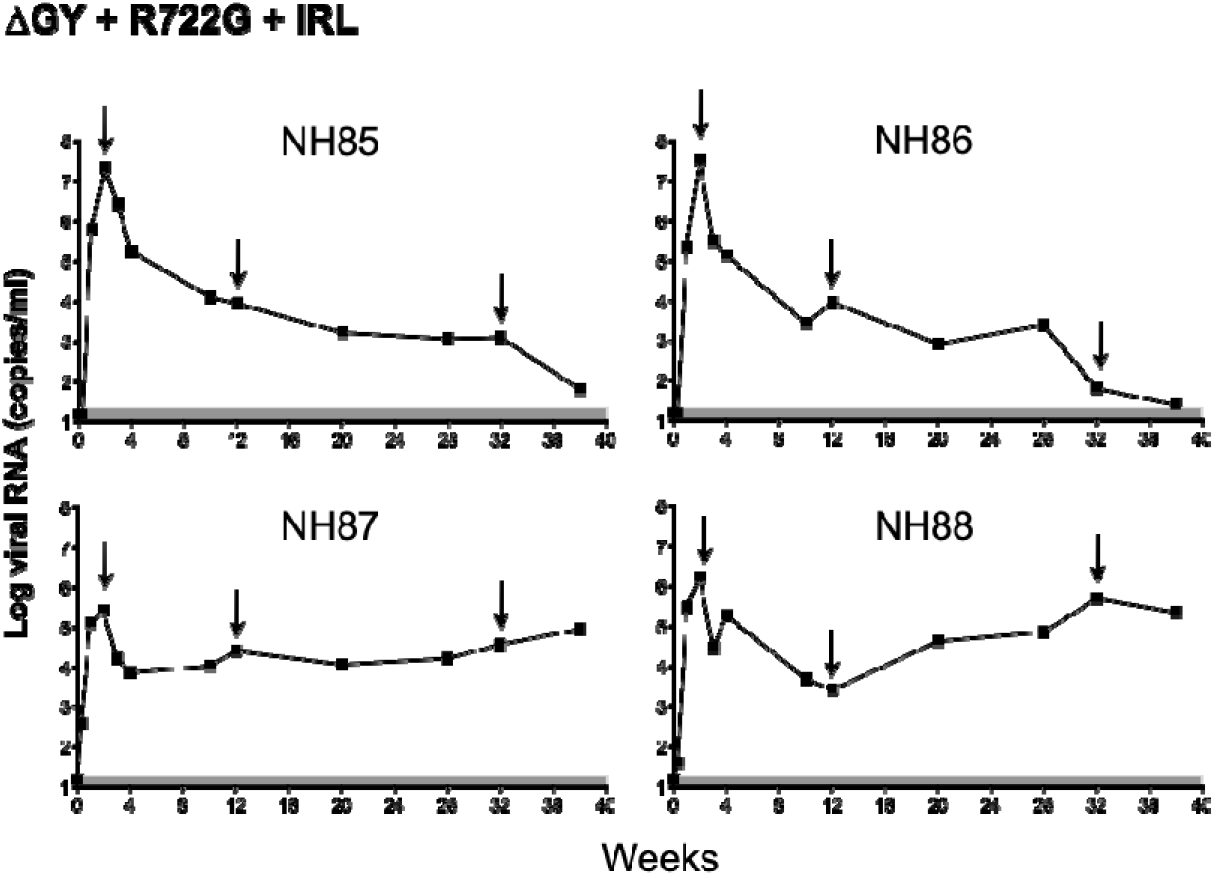
Plasma viral loads for pigtail macaques infected with SIVmac239ΔGY+R722G and the IRL set. Pigtail macaques were infected with SIVmac239ΔGY+R722G in combination with the IRL substitutions (T735I, Q739R, and P744L). Plasma viral loads are shown. Arrows denote time points at which plasma was obtained for SGA and sequencing. Animal identifiers are indicated.

SGA and sequencing of plasma virus was performed at multiple time points (Figure 8), and the results for the Env CD are shown for all amplicons (Supplemental Figure 10) with a summary of changes shown for a.a. 710-757 encompasing ΔGY, R722G, and the IRL set (Table 3), and a.a. 800-880 encompassing the distal CD (Supplemental Table 3). In 2 animals (NH85 and NH87) amplicons containing a G873E near the Env C terminus appeared at week 12 and increased to 100% of amplicons by week 33 (Supplemental Table 3 and Supplemental Figures 10 A and C). Interestingly, this mutation also evolved to a consensus mutation in KV73, the animal inoculated with SIVmac239ΔGY+R722G virus that initially developed the IRL substitutions (Supplemental Table 2 and Supplemental Figure 7F) and in a minority of amplicons in 2 of 3 animals infected with the SIVmac239ΔGY+R722G+ΔQTH (Supplemental Table 1 and Supplemental Figure 7). However, a G873R at this position appeared in NH86 and NH88 (in 100% and 93% of amplicons, respectively) suggesting that loss of G873 was likely driven by immune pressure rather than acquisition of a new trafficking signal. Additional mutations appeared in the CD and, for animal NH86, in the membrane spanning domain (Q712L in 19 of 24 amplicons at week 33), but none were common to all animals. As expected, all 4 animals developed R751G by week 12, consistent with ongoing replication of a SIVmac239-based virus (43). Remarkably, sequence analysis from weeks 2, 12 and 33 weeks post infection revealed conservation of all 3 IRL substitutions in nearly every amplicon at each time point, noteably maintaining P744L, which, as stated previously, generated a premature stop codon in the *tat* second exon (Table 3; Supplemental Figure 9).

**Table 3.**
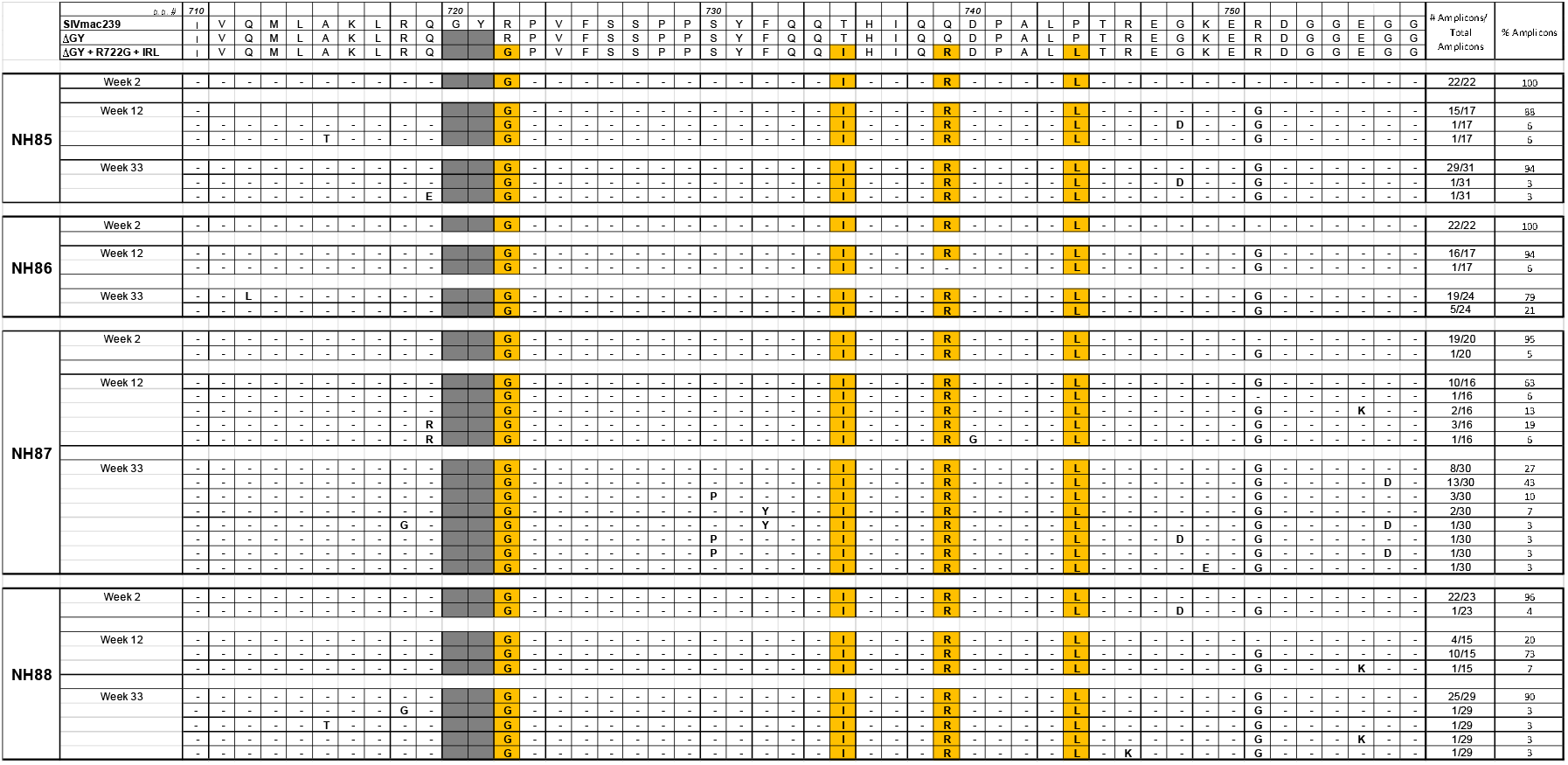
Viral evolution in macaques inoculated with SIVmac239ΔGY+R722G containing the IRL set. Summary of single genome amplification analysis of plasma virus performed at the indicated time points for 4 pigtail macaques inoculated with SIVmac239ΔGY+R722G containing the T735I, Q739R and P744L substitutions that arose in animal KV73 that progressed to AIDS (see Table 2). Sequences for SIVmac239,SIVmac ΔGY and SIVmac239ΔGY+R722G+IRL are shown at the top. Complete listings of a.a. sequences in the cytoplasmic domain for individual amplicons are shown in Supplemental Figure 10.

Collectively, these findings indicate that the novel IRL polarized sorting signal acquired during pathogenic evolution of a SIVmac239ΔGY+R722G virus *in vivo*, was completely conserved in *de novo* infections of naïve PTM. Although not sufficient to confer AIDS, at least through 33 weeks of infection, these findings indicate that acquisition of this novel gain-of-function signal for polarized sorting, but not endocytosis, was sufficient to restore high replicative capacity and fitness to a ΔGY+R722G virus. The finding that loss of the *tat* 2^nd^ exon, which was required to generate this signal, was retained throughout infection suggests that there was remarkably strong selection pressure *in vivo* to maintain the polarized sorting function of Env.

## DISCUSSION

The cytoplasmic domains of HIV and SIV Envs contain a common, highly conserved Tyr-based trafficking motif (GYxxØ) that mediates both clathrin-dependent endocytosis (17, 20–22) and polarized sorting (27, 39, 51). Though less studied, similar motifs have been described in the Env cytoplasmic domain of other retroviruses, including HTLV-1 (52). For HIV and SIV Envs, the GYxxØ motif can mediate endocytosis through interaction with the heterotetrameric clathrin adaptor protein AP2, analogous to the uptake of cellular proteins that contain similar Tyr-based YxxØ signals, with or without a proximal Gly (17, 20–22, 53). By contrast, the pathways and cellular partners required for polarized sorting of HIV and SIV Env are less well defined, but the Tyr within the GYxxØ motif is critical (25). Additional cellular motifs including di-leucine, [D/E]xxxL[L/I], DxxLL and acidic clusters, have also been implicated in both the endocytosis and polarized sorting of cellular proteins (37) as well as HIV-1 and SIV Env (22, 54). Although there are examples of how the loss of viral Env trafficking motifs can alter pathogenesis in small animal models (55, 56), for HIV and SIV the *in vivo* role of these signals has remained unclear. In this paper we provide the first demonstration that the membrane proximal GYxxØ in the cytoplasmic domain of SIV Env is crucial for SIV pathogenesis in rhesus and pigtail macaques and that the individual functions associated with this motif are under strong positive selective selection.

Several pathogenic roles have been proposed for trafficking functions in HIV and SIV Envs. With regard to endocytosis, because Env delivered to the cell surface is rapidly internalized through clathrin-mediated endocytosis, resulting in low steady state levels of Env on the plasma membrane of infected cells, we and others have proposed that Env internalization could contribute to immune evasion, rendering infected cells less susceptible to antibody attack either directly or by antibody-dependent cell-mediated cytotoxicity (17, 21, 57, 58). Consistent with an *in vivo* role for the GYRPV motif, SIVmac239 with a T721I substitution remained stable during extensive serial passaging *in vitro* but rapidly reverted in both RM and PTM (34 and unpublished observations). In contrast to point mutations in the HIV or SIV GYxxØ motif, the ΔGY mutation in SIVmac239 globally reduces Env content on cells and within virions (Figure 2). For HIV-1, substitutions of the Tyr within the analogous GYSPL consensus sequence are poorly tolerated even during *in vitro* replication (59), suggesting a role for this motif in assembly and/or infectivity, although this effect can vary with different viral strains (59, 60). In contrast to the role of endocytic function, the relevance of polarized sorting of HIV and SIV Env *in vivo* has been unclear, although this property has been recognized to positively affect viral infection and cell-cell spread *in vitro* (25–28).

We have shown that deleting Gly and Tyr from the GYRPV motif in SIVmac239, results in a novel phenotype *in vivo*. In the majority of PTM infected with SIVmac239ΔGY, acute viral replication occurs to near wildtype levels, but with the onset of host immune responses is well controlled through cellular immune responses but not neutralizing antibodies (18). In contrast to SIVmac239 and despite robust SIVmac239ΔGY replication in lymphoid tissues, there is only transient infection of gut CD4+ T cells, no detectable infection of macrophages, and little to no immune activation (18). However, progression to AIDS has been reported in SIVmac239ΔGY-infected RM (32) and rarely in PTM (18) where it occurred in association with novel changes in the Env CD. The role of these changes and the extent to which they compensate for defects introduced by the ΔGY deletion have been unclear (61).

In the current study we evaluated mutations acquired during pathogenic SIVmac239ΔGY infection, which included an R722G substitution flanking the ΔGY deletion and the loss of 3 amino acids (ΔQTH) resulting from a remarkable 9 nt deletion that occurs in overlaping reading frames and splice acceptor sites for the 2^nd^ exons of *rev* and *tat*. The ΔQTH deletions created novel YFQI or YFQL sequences in Env reminiscent of the YxxØ trafficking motifs of cellular proteins (Figure 1)(17). We showed that YFQI and YFQL restored both endocytic and polarized sorting functions of the parental SIVmac239 GYRPV motif. Reacquisition of these functions depended on the Tyr, and for endocytosis required AP2, indicating that viral use of cellular trafficking machinery was reestablished and correlated with progression to AIDS. We also showed that the ΔGY deletion reduced Env on virions, likely due to a general reduction in Env content within SIVmac239ΔGY-infected cells (Figure 2), and that this defect could be rescued by R722G but not ΔQTH. While R722G did not restore endocytic or polarized sorting functions, it was critical to maintaining replication fitness of ΔGY viruses containing ΔQTH deletions (Supplemental Figure 4). Interestingly, the R722G substitution regenerated a RQG sequence (a.a. 718-720) present in SIVmac239 Env CD. We also demonstrated that an S727P substitution, previously seen in SIVmac239ΔGY-infected RM (33) and one PTM (34) that progressed to AIDS, also restored Env content on ΔGY virions to wildtype levels, similar to the R722G substitution. In all ΔGY-infected animals that progressed to disease, either R722G or S727P appeared (although never together in the same transcript), suggesting that restoring Env expression levels was critical for progression. In an earlier study, we showed in 4 RM infected with SIVmac239ΔGY, S727P increased infection and depletion of gut CD4+ T cells (33). Of 2 animals followed through chronic infection, one progressed to AIDS, associated with appearance of a new YxxL motif (YTLL) in the Env CD, and the other controlled virus to undetectable levels without additional changes, similar to the SIVmac239ΔGY+R722G infected KV75 in the current study (Figure 7; Supplemental Figure 5A). Although the trafficking functions of the YTLL sequence were not analysed, the data suggested that while increasing Env content in cells and/or on virions (by either S727P or R722G) is required for pathogenicity, it is not sufficient, and that a gain of Env trafficking functions is also required.

To examine the role of these novel changes on SIV infection *in vivo*, we infected PTM with SIVmac239ΔGY containing mutations encoding R722G and the ΔQTH deletion. Significantly, the mutations persisted in all amplicons and at all time points (Supplemental Figure 7). In contrast to SIVmac239ΔGY-infected PTM, where viral loads are typically controlled to low or undetectable levels (18), partial reconstitution of SIVmac239 pathogenicity was observed with one animal (KV74) progressing to AIDS, and detectable viremia persisted in all 3 animals for up to 64 weeks (Figure 6). When PTM were infected with SIVmac239ΔGY containing R722G, 2 of 3 animals progressed to AIDS, with high viral loads associated with the appearance of additional changes in the Env CD, either a new ΔQTH deletion generating a YFQL (Table 2 Supplemental Figure 7E) or 3 substitutions (T735I, Q739R, and P744L) (Table 2 Supplemental Figure 6F). This latter IRL set generated a novel signal for polarized sorting, but not endocytosis, and also created a stop codon within the *tat* 2^nd^ exon associated with P744L, which deleted the C-terminal 22 a.a. of Tat. When a SIVmac239ΔGY+R722G virus containing IRL was given to 4 PTM, all animals maintained persistent viremia to 30 weeks and, notably, the IRL substitutions were highly conserved through 33 weeks of infection.

Our findings that the ΔQTH deletion or IRL substitutions restored Env endocytosis and polarized sorting and were retained during *de novo* infections, indicates that these trafficking functions are likely critical for pathogenic SIV infection. While AP2-mediated Env endocytosis can be directed by the membrane proximal GYxxØ motif and membrane distal LL-like motifs (21, 22, 54, 62, 63) experiments with short and long cytoplasmic domain constructs in MDCK cells indicate that the GYxxØ motif is the only determinant in the SIV Env cytoplasmic domain mediating polarized sorting (Figure 5). The finding that the IRL substitutions restored the basolateral sorting function lost by the ΔGY mutation but not endocytosis (Figure 7; Supplemental Figure 2), and was sufficient to confer persistent viremia to a SIVmac239ΔGY+R722G virus, suggests that polarized sorting of Env may be the principal trafficking function of the GYxxØ motif during progression to disease. This is not to say that Env endocytosis is not important, rather that this function may be carried out by less well defined endocytic signals present in SIV Env cytoplasmic domain (22) similar to the highly conserved C-terminal di-leucine that we have shown mediates HIV-1 Env endocytosis (54). Nevertheless, for the 4 animals given SIVmac239ΔGY+R722G containing the IRL substitutions, this additional polarized sorting function imparted higher and more sustained viral loads.

The mutations generating ΔQTH and the IRL set arose in an area of the genome in which all three reading frames are used, with the potential to affect other viral transcripts. Notwithstanding, the impact of the gain-of-function mutations in Env on Tat and Rev were extraordinary and included, for ΔQTH, deletion and point mutations and loss of a splice acceptor site (Supplemental Figure 8), and, for IRL, premature truncation of the *tat* 2^nd^ exon (Supplemental Figure 7). Although for SIV and HIV the *tat* 2^nd^ exon can be dispensible for viral replication *in vitro* (64, 65), in RM infected with SIVmac239 lacking a *tat* 2^nd^ exon, reversion to a two exon *tat* occurred in association with high viral loads and falling CD4+ T cell counts, whereas persistence of a single exon *tat* was associated with viral control (66). It is therefore remarkable that the P744L substitution, which created the *tat* stop codon during evolution of the IRL substitutions, was maintained when SIVmac239ΔGY containing these mutations was used to infect naïve animals. Thus, as the IRL signal for polarized Env trafficking was maintained, the *tat* 2^nd^ exon proved to be dispensible in our study. These observations are remarkable as SIVmac239ΔGY containing the IRL mutations contains two known potent fitness costs (GY and P744L) where ΔGY alone is sufficient to potently control viral replication. These findings further emphasis the strong selection pressure *in vivo* to generate and maintain Env trafficking functions.

Unlike epithelial cells, which maintain fixed apical and basolateral plasma membrane domains, T cells undergo polarization during their migration along chemokine gradients, (67–71), during the formation of immunological synapses with antigen presenting cells or the cellular targets of cytotoxic T cells (72), and when virological synapses form between HIV- and SIV-infected T cells and uninfected T cells and macrophages (73). Virological synapses require interactions between Env and CD4 (74–78) that ultimately enhance the efficiency of viral infection and cell-cell spread *in vitro* (25, 76, 79, 80). Although HIV Gag and RNA colocalize at the uropod of polarized T cells in an Env-independent manner (29, 31, 81), Env is also present at this site, indicating that, similar to murine leukemia virus (82) and measles virus (55), the uropod is a site for viral assembly as well as engaging CD4 on target cells to nucleate synapse formation (76, 79). It is likely that polarized trafficking of Env contributes to this process. While virologic synapses and polarized sorting functions of Env have been recognized as contributing to pathogenesis of MuLV (56, 82) and measles virus (55), there is little direct evidence that this function is important for HIV or SIV *in vivo*. Our findings that the ΔQTH and IRL mutation set, acquired in SIVmac239ΔGY-infected animals that progressed to AIDS, recovered endocytic and/or polarized sorting signals and were retained during *de novo* infections, indicates that these trafficking functions are likely critical for SIV replication and pathogenesis.

In summary, our characterization of pathological revertants in SIVmac239ΔGY-infected macaques highlight critical *in vivo* roles played by cellular trafficking motifs in the SIV and, by analogy, HIV-1 Env cytoplasmic domains. Whereas the ΔGY deletion within the conserved GYxxØ motif ablates signals for endocytosis and polarized sorting, these functions were regained through novel deletions or substitutions at the expense of collateral changes in Tat and Rev. Given that SIVmac239ΔGY replicates poorly in gut CD4+ T cells and fails to infect macrophages *in vivo* (18, 32), our findings suggest that trafficking functions, particularly the polarized sorting of Env, could be required for optimal infection of these cells, perhaps by promoting virological synapse formation and viral spreading through cell-cell contacts. Virologic synapses enhance the efficiency of viral infection and cell-cell spread *in vitro* (25, 79), in part at least through the generation of transcriptional signals (80), and have been shown to enable viruses to overcome diverse barriers to infection, including restriction factors (83, 84), neutralizing antibodies (76, 85) and reduced levels of receptors on target cells (86). SIVmac239 infection of macrophages, which have low levels of CD4, has been shown to require cell-cell transmission (77) and could be compromised viruses with Envs lacking the ability to form these structures. Consistent with the view that T cell/macrophage interactions are compromised during ΔGY infection, comparisons of transcriptional profiles between ΔGY and SIVmac239-infected macaques have recently shown the striking absence of macrophage transcripts during acute ΔGYinfection (manuscript in preparation) in marked contrast to SIVmac239 infection where monocyte/macrophage transcripts are highly upregulated (7). Studies are ongoing in SIVmac239ΔGY and SIVmac239-infected macaques to assess directly virologic synapse formation *in vivo* as well as the contributing role of compensatory mutations that may promote cell-to-cell spread.

## MATERIALS AND METHODS

### Ethics statement

Pigtail macaques used in this study were purpose bred at either the University of Washington National Primate Research Center or Johns Hopkins and moved to Tulane for these experiments. Macaques were housed in compliance with the NRC Guide for the Care and Use of Laboratory Animals and the Animal Welfare Act. Animal experiments were approved by the Institutional Animal Care and Use Committee of Tulane University (protocols P0088R, P0147, and P0312). The Tulane National Primate Research Center (TNPRC) is fully accredited by AAALAC International (Association for the Assessment and Accreditation of Laboratory Animal Care), Animal Welfare Assurance No. A3180-01. Animals were socially housed, indoors in climate-controlled conditions with a 12/12-light/dark cycle. All the animals on this study were monitored twice daily to ensure their welfare. Any abnormalities, including those of appetite, stool, behavior, were recorded and reported to a veterinarian. The animals were fed commercially prepared monkey chow twice daily. Supplemental foods were provided in the form of fruit, vegetables, and foraging treats as part of the TNPRC environmental enrichment program. Water was available at all times through an automatic watering system. The TNPRC environmental enrichment program is reviewed and approved by the IACUC semiannually. Veterinarians at the TNPRC Division of Veterinary Medicine have established procedures to minimize pain and distress through several means. Monkeys were anesthetized with ketamine-HCl (10 mg/kg) or tiletamine/zolazepam (6 mg/kg) prior to all procedures. Preemptive and post procedural analgesia (buprenorphine 0.01 mg/kg or buprenorphine sustained-release 0.2 mg/kg SQ) was required for procedures that would likely cause more than momentary pain or distress in humans undergoing the same procedures. The above listed anesthetics and analgesics were used to minimize pain or distress associated with this study in accordance with the recommendations of the Weatherall Report. The animals were euthanized at the end of the study using methods consistent with recommendations of the American Veterinary Medical Association (AVMA) Panel on euthanasia and per the recommendations of the IACUC. Specifically, the animals were anesthetized with tiletamine/zolazepam (8 mg/kg IM) and given buprenorphine (0.01 mg/kg IM) followed by an overdose of pentobarbital sodium. Death was confirmed by auscultation of the heart and pupillary dilation. The TNPRC policy for early euthanasia/humane endpoint was included in the protocol in case those circumstances arose.

### Antibodies, Reagents and Cell Lines

#### Antibodies

The following reagents were obtained from the Centre for AIDS Reagents National Institute for Biological Standards and Control [NIBSC], South Mimms, UK): Anti-Gag p57/27 (SIV 27c donated by P. Szawlowski or KK60 donated by K. Kent), anti-Nef (KK77 donated by Dr K. Kent), anti-CD4 (Q4120, provided by Q. Sattentau, University of Oxford). Murine anti-envelope monoclonal antibodies DA6 (to SIVmac gp120/gp160) and 35C11 (to SIVmac gp41) have been previously described (87). Anti-Gag p27 (3A8 was provided by J. McClure). Anti-CD3 (SP34), CD4 (L200) and CD8 (SK1 or SK2) were used for flow cytometry and obtained from BD Biosciences.

#### Reagents

Human sCD4 domains 1-4 (ARP6000 Progenics Pharmaceuticals, Inc, USA was obtained from the Centre for AIDS Reagents, NIBSC) and human sCD4 domains 1-2 (#7356 from Pharmacia, Inc was obtained through the NIH AIDS Reagent Program, Division of AIDS, NIAID, NIH).

#### Cell Lines

human embryonic kidney cells (HEK-293T; ATCC, CRL-3216), rhesus macaque kidney cells (LLC-MK2; ATCC, CCL-7), HeLa cells (from D. Cutler, MRC LMCB, UCL) and Madin Darby canine kindey cells (MDCKII from G. van Meer, University of Utrecht) were maintained in DMEM supplemented with 10% fetal calf serum (FCS; Sigma A7906) and 100 U/ml Pen/100 μg/ml Strep (Gibco, 15140-122). All cell lines used in this study were monitored for mycoplasma infection using the MycoAlert Mycoplasma Detection Kit (Lonza TL07-218).

### VSV-G pseudotyping of SIVmac239 and virus titration

HEK293T cells were co-transfected with full-length SIVmac239 genome constructs and a plasmid encoding the VSV-G glycoprotein (pMD2.G; provided by P. Mlcochova, Div. Infection and Immunity, UCL), at a ratio of 3 μg of provirus to 1 μg of pMD2.G, using the Fugene6 transfection reagent (Promega). Virions were concentrated from the culture medium 48 hours post infection (hpi) by first clearing large debris (centrifugation for 5 min at 2000 rpm) followed by ultracentrifugation through a 20% (w/v) sucrose cushion (23,000 rpm [98,000 × g], 2 hr, 4°C). The pellet was suspended in culture medium (RPMI-1640, 100 U/ml Pen/100 μg/ml Strep and 10% FCS) and stored under vapor phase of liquid nitrogen. Infectious titres were measured by serial dilution on LLC-MK2 cells; following infection for 48 hr, cells were washed in PBS pH 7.4, fixed (3% [w/v] formaldehyde, PBS pH 7.4) for 30 min at 4°C and stored overnight (0.1% [w/v] formaldehyde, PBS pH 7.4). The following day, samples were quenched (50 mM NH_4_Cl, PBS pH 7.4) for 15 min at room temperature (RT), permeabilised (0.1% [w/v] saponin, 1% [v/v] FCS, PBS pH 7.4) and immunolabeled with primary antibodies (anti-Gag [SIV 27c] and/or anti-Nef [KK77]) for 1.5 hr at RT followed by secondary antibodies conjugated to Alexa Fluor dyes (Invitrogen) for 1 hr at RT. Viral protein expression was detected by imaging with Opera LX and Phenix high content imaging platforms (Perkin Elmer) and the number of infected cells determined using a Columbus Analysis system (Perkin Elmer).

### Biochemical analysis of viral protein expression by western blotting

LLC-MK2 cells were incubated with VSV-G pseudotyped SIVmac239 to infect 40% of the cell population. The cell culture medium was replaced at 24 hpi and the cells cultured for a further 48 hrs. At 72 hpi, viruses in the cuture supernatants were recovered by centrifugation through sucrose, as described above, and the corresponding cells were lysed in 150 mM NaCl, 1% (v/v) Triton, 50 mM Tris/HCl pH 8.0 containing complete protease inhibitors (Roche). The lysates were cleared of insoluble material and stored at −80°C prior to analysis. Cell and viral lysates were mixed with Laemmli Sample Buffer containing 100 mM dithiothreitol (DTT) and heated for 10 min at 98°C. To enable clear separation of Env and Gag proteins on the same gel, proteins were separated on Laemmli SDS-polyacrylamide gels where the resolving gel comprised an upper gel of 8% acrylamide over an equal volume of 15% acrylamide. Following electrophoresis, the proteins were electroeluted from gels onto Immobilon-F PVDF membranes (Millipore) at 100 mA for 16-18 hr at 4°C under wet blotting conditions (10 mM 3-[Cyclohexylamino]-1-propanesulfonic acid pH 11.0, 10% [v/v] methanol). Membranes were blocked (5% [w/v] non-fat dried milk in 0.1% [v/v] Tween-20, PBS pH 7.4) for 3 h, followed by detection of proteins with primary antibodies (0.5% [v/v] Tween-20/ 1% [w/v] Bovine Serum Albumin [BSA], PBS pH 7.4) for 1.5 hr at RT, or 4°C overnight, followed by secondary antibodies conjugated to IRDye 800CW or IRDye 680 (LiCOR Biosciences) for 1.5 hr at RT. Viral proteins were detected with antibodies to gp120/gp160 (DA6), gp41 (35C11) and Gag p57/p27 (KK60). Images of the western blots and the intensity of protein bands were obtained and quantified using an Odyssey infrared imaging system (LiCOR Biosciences).

### Biochemical analysis of cell surface protein levels

At day 3 post infection, LLC-MK2 cells (40% infected) were washed with ice-cold PBS and cell surface proteins covalently labelled with cell impermeable EZ-Link Sulfo-NHS-S-S-Biotin (0.5 mg/ml; Pierce) for 45 mins at 4°C. Excess label was removed and the samples quenched by washing with TBS (154 mM NaCl, 10 mM Tris/HCl pH 7.4) at 4°C. Cell lysates were prepared as described above and diluted to equal protein concentrations. An aliquot of each lysate (150 μg of protein) was incubated with 100 μl, 50% slurry of NeutrAvidin Agarose beads (Pierce) overnight at 4°C with inversion. To show that all the biotinylated proteins were captured with this first incubation, the lysate was separated from the beads and the process repeated with fresh NeutrAvidin beads for 3 hr. Subsequently, the beads were washed once with lysis buffer, once with TBS and once with TE (10 mM Tris/HCl pH 7.4 and 5 mM EDTA) and eluted twice by incubation with Laemmli sample buffer, containing 100 mM DTT, and heating for 10 min at 98°C. Cell lysates (‘L’; equivalent to 30 μg of protein) were separated alongside the proteins eluted from the NeutrAvidin beads (‘S’; Surface) on SDS-PAGE gels, as described above. Following electrophoresis, the proteins were transferred to Immobilon-F PVDF membranes (Millipore) at 0.8 mA/cm^2^ for 2 hr at RT under semi-dry blotting conditions using a discontinuous 3 buffer system (Anode Buffer I: 0.3 M Tris pH 10.4, 10% MeOH; Anode Buffer II: 25 mM Tris pH 10.4, 10% MeOH; Cathode Buffer: 25 mM Tris, 40 mM 6-amino-n-caproic acid pH 10.4) modified from (88). Proteins were detected and imaged as described above.

### *In vitro* replication of SIV isolates in rhesus and pigtail macaque PBMCs

Purified peripheral blood mononuclear cells (PBMCs) from rhesus or pigtail macaques stored at −140°C were thawed and cultured for 72 hours in RPMI with 5 μg/ml concanavalin A (Sigma-Aldrich) at a concentration of 2-3 x 10^6^ cells/ml. After 72 hr, cells were washed and resuspended 1×10^6^ cells/ml in RPMI with 100 U/ml rHu IL-2 (Aldesleukin, Prometheus Laboratories, Inc.) and infected with viruses (250 ng of p27 Gag). After 24 hr, cells were washed and cultured in fresh RPMI-complete medium supplemented with IL-2 and the supernatant sampled for reverse transcriptase activity at 0, 3, 6, 10 and 14 days post inoculation, as described (89).

### Biochemical analysis of virion envelope content from primary macaque PBMCs

Rhesus macaque PBMCs were infected with viral isolates as for replication assays above. Six days post-inoculation, cell-free supernatant was removed and virions pelleted through 20% sucrose by ultracentrifugation for 120 minutes. Viral pellets were resuspended in 1X TNE buffer, and quantified by p27 ELISA. The samples were reduced by combining NuPAGE 10X Reducing Agent (Life Technologies) and NuPAGE 4X LDS Sample Buffer (Life Technologies) and incubating at 95°C for 10 mins. Equal amounts of p27 were loaded for PAGE and proteins transferred to a PVDF membrane and blocked with 5% NFM for 1 hr at RT. Membranes were cut in half and blotted separately with murine anti-Gag (3A8) or anti-gp120 (DA6) antibodies. Blots were incubated with goat anti-mouse horse radish peroxidase (HRP)-conjugated secondary antibody and the bands developed with Luminata Forte Western HRP Substrate (Merck Millipore).

### Stable cell lines

HeLa cells were transfected with plasmids encoding CD4-SIV Env chimeras (22) using TransIT-HeLaMONSTER (Mirus) and stable transfectants were selected with 400 μg/ml G418 Sulphate (Calbiochem). The stable transfectants were enriched for the expression of CD4 by FACS. Briefly, transfected HeLa cells were detached with PBS-EDTA (PBS, 5 mM EDTA, pH 7.4), washed, resuspended in ice-cold RPMI-1640 containing 1% (v/v) FBS and incubated with 5 μg/ml anti-CD4 (Q4120) for 1 hr. Subsequently, the cells were washed 3 times with RPMI/1% FCS, to remove unbound antibody, and then incubated with 2 μg/ml Alexa Fluor-488 conjugated secondary antibody for 1 hr at 4°C. Finally, cells were washed once with RPMI/1% FCS, twice with PBS/1% FCS and sorted by FACS (FACSAria, Becton Dickinson). MDCKII cells were transfected by electroporation with plasmids encoding CD4-SIV Env chimeras using an Amaxa Cell Line Nucleofector™ Kit L and Amaxa Nucleofector II with settings L-05 (Lonza). Stable transfectants were selected with 400 μg/ml G418 Sulphate (Calbiochem). Alternatively, stable MDCKII cells were generated by transfection with HIV-based pseudoviruses encoding SIVmac239 Env constructs or CD4-SIV Env chimeras as controls. Briefly, VLPs were generated in HEK 293T cells by co-transfection of pCMV-d8.91, encoding HIV-1 *gag-pol* (enabling the formation of viral cores) together with the coding sequence for SIVmac239 Env cloned into the dual promoter self-inactivating vector pHRSIN-CSGWdINotI_pUb_Em, (90, 91), and VSV-G (pMD2.G), using Fugene 6 (Promega).

### siRNA knockdowns

HeLa cells (1.67×10^6^ per 10 cm dish) expressing CD4-SIV Env chimeras were seeded and transfected the same day with siRNA oligonucleotides (145 nM) using Oligofectamine (Invitrogen). Oligonucleotides targeting the μ2 subunit of adaptor protein complex 2 (AP2) were used as duplexes with 3’-dTdT overhangs as previously described (38) (RNA sequences: sense, 5’ GAUCAAGCGCAUGGCAGGCAU; antisense, 5’ AUGCCUGCCAUGCGCUUGAUC [Dharmacon]). Cells were passaged and used for endocytosis assays and western blotting 24 hr later. To assess the knockdown efficiency, cells were lysed as described above, cleared of insoluble material and 30 μg cell protein separated by Laemmli SDS-PAGE. Gels were electroeluted onto Nitrocellulose (BioTrace NT, Pall) 1.2 mA/cm^2^ for 1 hr at RT under semi-dry blotting conditions (25 mM Tris, 192 mM glycine, 0.1% [w/v] SDS, 20% [v/v] MeOH). Proteins were detected and imaged as described above.

### Endocytosis Assay

Quantitative endocytosis assays were performed using HeLa cells expressing CD4-SIV Env chimeras. Cells (42×10^3^ cells/well in 4 or 24 well plates) were seeded 24 hr prior to use. For analysis, the cells were rapidly cooled with ice-cold media (RPMI-1640, 20 mM HEPES, 10 mM Bicarbonate and 0.2% [w/v] BSA, pH 7.0) and incubated with an anti-CD4 antibody (5 μg/ml Q4120) for 1 hr at 4°C. Unbound antibody was washed away and endocytosis was initiated by rapidly warming with 37°C media and subsequently stopped by rapidly cooling with 4°C media at the indicated times. The anti-CD4 remaining at the cell surface was detected with HRP-conjugated anti-mouse IgG. Subsequently, cells were washed once with ice-cold media, twice with PBS, lysed (150 mM NaCl, 1% (v/v) Triton, 10 mM Hepes pH 7.0, and complete protease inhibitors [Roche]), and cleared of insoluble material. HRP in the cell lysate was detected by the addition of 50 μM Amplex Red (Invitrogen, in 150 mM NaCl, 1% [v/v] Triton, 200 μM H_2_0_2_, 10 mM HEPES pH 7.0) and measuring the rate of production of resorufin (Δfluorescence/min where the reaction obeys first order kinetics) with an EnVision Multilabel Reader (Perkin Elmer).

### Microscopy

HeLa cells were seeded on 13 mm, #1.5, glass coverslips 24 hr prior to use. Cells were incubated with 10 μg/ml anti-CD4 (Q4120) in media (DMEM, 1% FBS) for 3 hr at 37°C or left untreated. Cells were washed (PBS pH 7.4 containing 0.9 mM CaCl_2_ and 0.49 mM MgCl_2_ [PBS^++^]), fixed (PBS^++^ containing 3% [w/v] formaldehyde) for 20 min, quenched (50 mM NH_4_Cl in PBS pH 7.4) for 15 min, permeabilized (0.05% [w/v] saponin, 1% [v/v] FCS in PBS pH 7.4) and immunolabeled with 10 μg/ ml anti-CD4 (Q4120) for 1.5 hr at RT followed by detection with secondary antibodies conjugated to Alexa Fluor dyes (Invitrogen). Finally, samples were washed with water and mounted on coverslips with Mowiol 4-88 (Calbiochem). Images were acquired using Nyquist criterion with a Leica TCS SPE confocal system with a 63x ACS APO/ NA 1.3 oil immersion lens and galvanometer driven stage insert and deconvolved using Huygens software.

MDCKII cells (2.7×10^5^/cm^2^) were seeded on polyester Transwell 0.4 μm clear filters (Corning) and cultured for 6 days to establish polarized monolayers (trans-epithelial resistance of 80-100 Ω/cm2). Cells expressing CD4-SIV Env chimeras were washed twice (PBS^++^), fixed and quenched as described above, and immunolabeled with 10 μg/ ml anti-CD4 (Q4120) for 1.5 hr at RT. Cells expressing SIVmac239 envelope protein were rapidly cooled with ice-cold media (DMEM, 1% [v/v] FBS), incubated with 480 nM human sCD4 (either D1-4 ARP6000 or D1-2 #7356) for 15 min prior to the addition of 10 μg/ml of anti-Env monoclonal (7D3). After 1 hr, the cells were washed, fixed (PBS^++^ containing 3% [w/v] formaldehyde) for 30 min 4°C and quenched (50 mM NH_4_Cl in PBS pH 7.4) for 15 min at RT. All samples were permeabilized (0.05% [w/v] saponin, 1% [v/v] FCS in PBS^++^) and primary antibodies were detected with secondary antibodies conjugated to Alexa Fluor dyes (Invitrogen). Images were acquired as above. To identify the fluorescent signal from the lateral and apical membranes, cells were co-stained or E-cadherin (lateral) and EZ-Link Sulfo-NHS-S-S-Biotin (Pierce, as described above) added apically and detected with streptavidin Cy5 (Jackson). In addition, apical and basolateral membranes were also defined for CD4-SIV Env ΔGY where, after incubation with anti-CD4 antibody as described above, the membranes were immunostained using anti-mouse conjugated to a different colour Alexa Fluor dye (488 added to the basolateral side and 546 added to the apical side [not shown]). Identical boxes were drawn around the basolateral and apical membrane domains using ImageJ and the ratio of basolateral to apical labelling density calculated. For display purposes images were deconvolved using Huygens software.

### Animals, viral inoculations, and sample collection

A total of 10 pigtail macaques were used in this study and were inoculated intravenously (i.v.) with 100 50% tissue culture infective dose (TCID50) of SIVmac239ΔGY+R722G (n=3) or SIVmac239ΔGY+R722G+ΔQTH (n=3), or SIVmac239ΔGY+R722G+IRL (n=4). All animals were MHC genotyped and haplotypes are shown in supplemental table 4. Before any procedure, the animals were anesthetized by intramuscular injection of ketamine hydrochloride (10 mg/kg). Viruses were produced in HEK 293T cells transfected with plasmids containing full-length proviral DNA. Viruses were quantified by determining (TCID_50_) on rhesus macaque PBMCs. Groups of pigtail macaques housed at TNPRC, infected with either SIVmac239ΔGY or SIVmac239 and described in a previous study (18), were used for comparison. Prior to use, all animals tested negative for antibodies to SIV, simian T cell leukemia virus (STLV), and type D retrovirus and by PCR for type D retrovirus. Multiple blood samples and small intestinal biopsy samples (endoscopic duodenal pinch biopsy samples or jejunal resection biopsy samples) were collected under anesthesia (ketamine hydrochloride or isoflurane) at various times from each animal. Animals were euthanized if they exhibited a loss of more than 25% of maximum body weight, anorexia for more than 4 days, or major organ failure or medical conditions unresponsive to treatment (e.g., severe pneumonia or diarrhea) at the discretion of veterinarians.

### Quantitation of viral load in plasma

Plasma viral loads were determined at various times using a reverse transcription-PCR (RT-PCR) assay with a limit of detection of between 15 and 21 SIV RNA copies/ml (92).

### Lymphocyte isolation from intestinal tissues CD4 T cells in gut LPL

Intestinal cells were collected by endoscopic pinch biopsies of the small intestine from animals at various times. Intestinal biopsy procedures and isolation of cells from intestinal tissues were described previously (93). Intestinal cells were isolated using EDTA-collagenase digestion and Percoll density gradient centrifugation.

### SGA analysis

Single genome amplification (SGA) and Sanger sequencing was performed on plasma samples from infected PTM at various time points after infection. The entire env gene was sequenced using a limiting-dilution PCR to ensure that only one amplifiable molecule was present in each reaction mixture, as described (32, 94). Sequence alignments were generated with Geneious and presented as highlighter plots (www.hiv.lanl.gov) or alignment tables generated in DIVEIN (95). APOBEC signature mutations were identified with Hyper-Mut (www.hiv.lanl.gov).

## ACKNOWLEDGEMENTS

We thank Drs Catherine Hogan (Cardiff University) and Karl Matlin (University of Chicago) for advice and protocols for working with MDCK cells and Graham Warren (UCL) for critical comments on the manuscript. The project has been supported by the following funding: pigtail macaques acquired from Johns Hopkins were supported by NIH grant 5U42OD013117 prior to sale to Tulane; by Federal funds from the National Cancer Institute, National Institutes of Health, under Contract Nos. HHSN261200800001E and 75N91019D00024; by R01 AI138782-01 to JAH and by UK Medical Research Council funding to the MRC-UCL Laboratory for Molecular Cell Biology University Unit (MC_UU00012/1 and MC_U12266B) to MM.

## Supplemental Figures

**Supplemental Figure 1.**
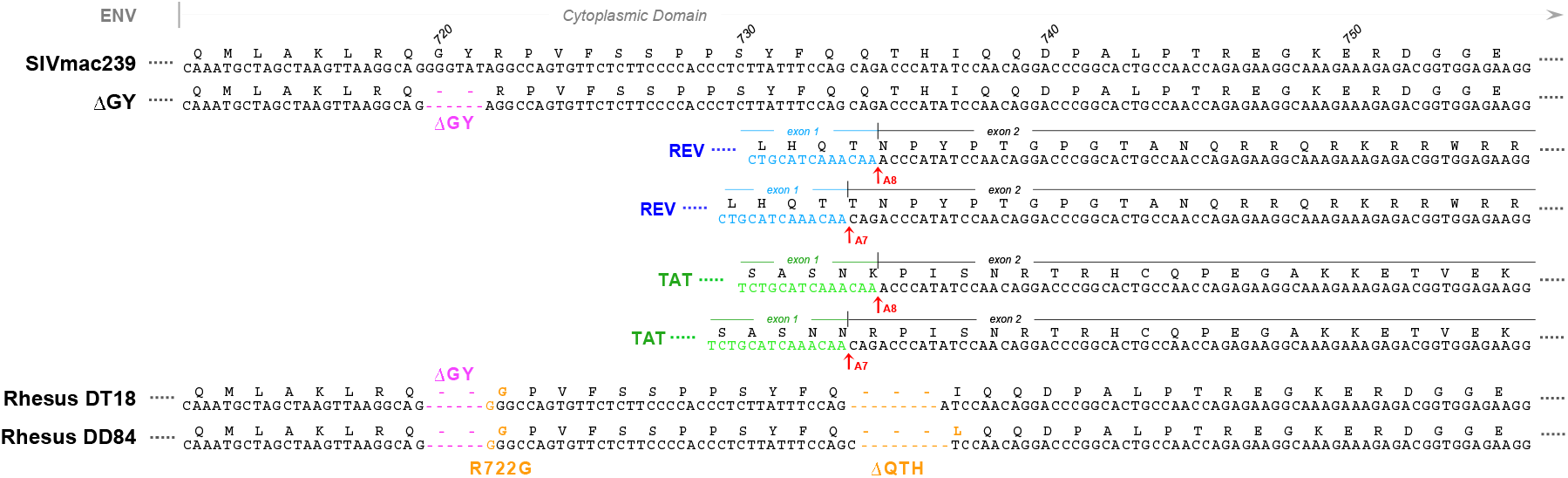
Effects of the acquired ΔQTH mutations on env, rev and tat open reading frames. Top Panel shows amino acid (a.a.) and nucleotide (nt) sequences for SIVmac239 and ΔGY Env, Tat and Rev proteins and mRNAs aligned by sequences in the Env CT. Known splice acceptor sites A7 and A8 are indicated for Rev and Tat with partial a.a. and nt sequences shown for the 1st exons of these proteins in blue and green, respectively. Bottom Panels show the deletion of QTH in Env (ΔQTH) that occurred in two rhesus macaques (RM) infected with SIVmac239ΔGY that progressed to AIDS (32). In RM DT18 ΔQTH resulted from loss of nt 8803-8811 and generated a new YFQI sequence in Env; in RM DD84 ΔQTH resulted from loss of nt 8804-8812 and generated a new YFQL sequence. Both ΔQTH mutations occur in regions of splice acceptor sites utilized for second exons of *rev* and *tat*. The effects of these mutations on Rev and Tat mRNA were subsequently determined *in vitro* and *in vivo* and are shown in Supplemental Figure 8. Both ΔQTH mutations arose in association with the R722G mutation shown flanking the ΔGY mutation site.

**Supplemental Figure 2.**
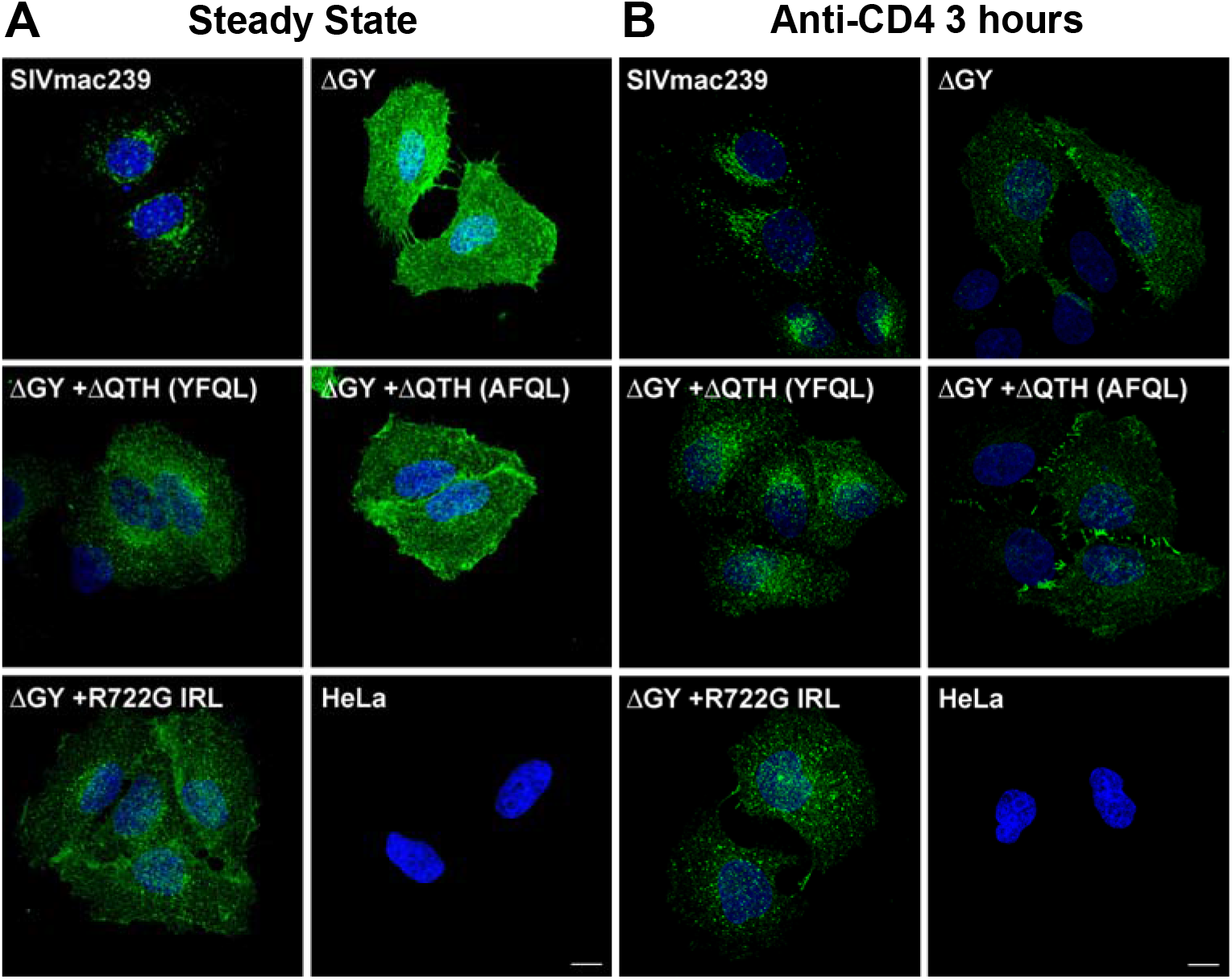
The ΔGY mutation and mutations acquired *in vivo* modulate cellular distribution and trafficking. **(A)** Steady state cellular distribution of CD4-SIV Env CD chimeras in HeLa cells (described in Figure 3, 5 and 7). **(B)** Cellular distribution of CD4-SIV Env CD chimeras in HeLa cells after incubation with anti-CD4 (Q4120) at 37°C for 3 hrs prior to fixation. Confocal Z stacks were deconvolved and displayed as maximum projections. Scale bar = 10 μm.

**Supplemental Figure 3.**
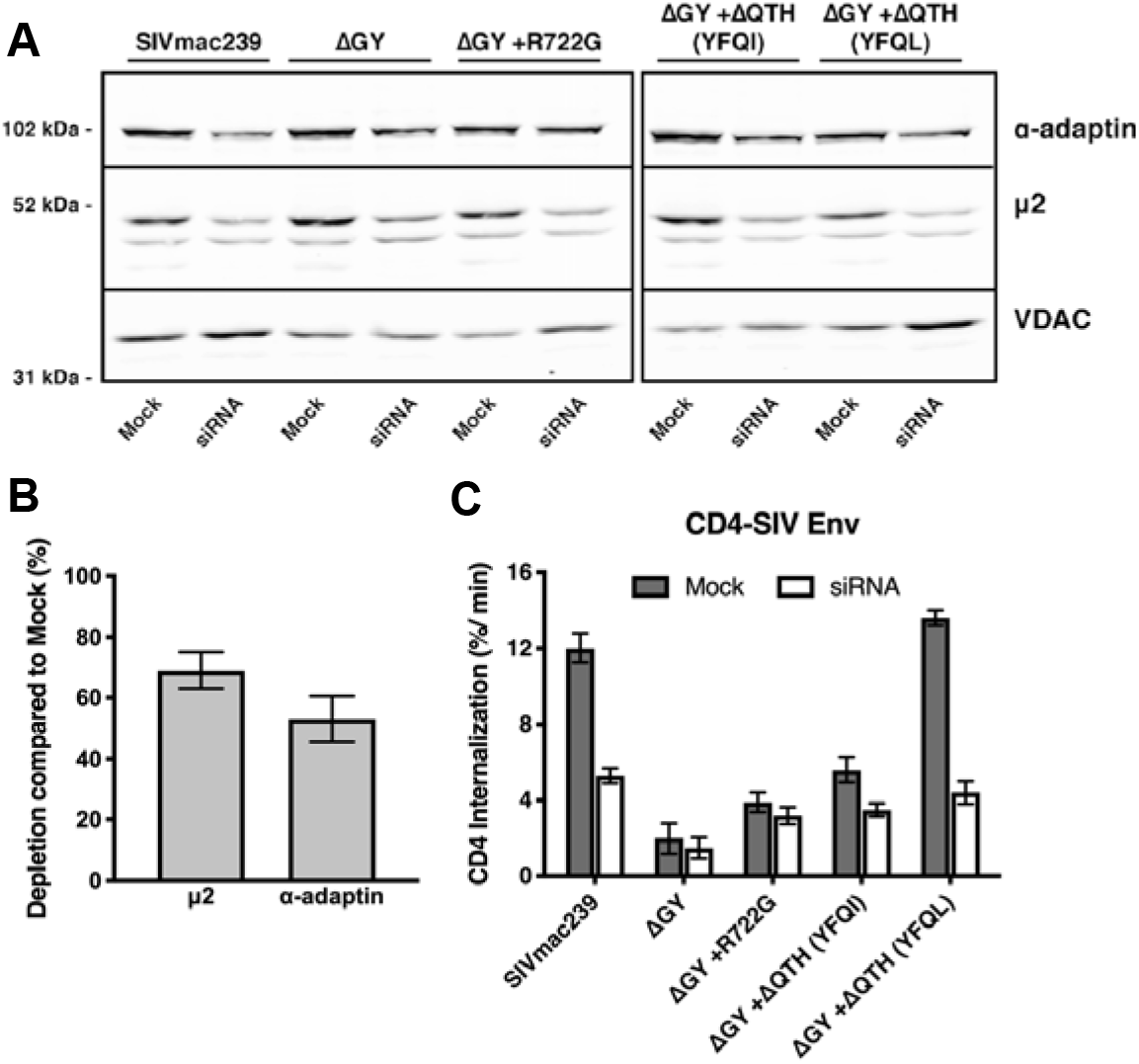
The ΔQTH mutations are AP2 dependent endocytic motifs. HeLa cells expressing CD4-SIV Env CD chimeras (Figure 3) were transfected with siRNA targeting the μ2 subunit of adaptor-related protein complex 2 (AP2). **(A)** A representative western blot of cell lysates incubated with antibodies to AP2 subunits (□adaptin or μ2) or with a loading control (anti-VDAC). **(B)** Quantitation of western blots. AP2 subunit depletion was calculated by comparison to mock conditions for all cell lines. **(C)** Endocytic rates of CD4-SIV Env short tail constructs +/- μ2 siRNA transfection. Results are expressed as the rate of CD4 endocytosis over the first 5 minutes after warm up. Graphs show the mean ± SEM calculated from n≥4 independent experiments.

**Supplemental Figure 4.**
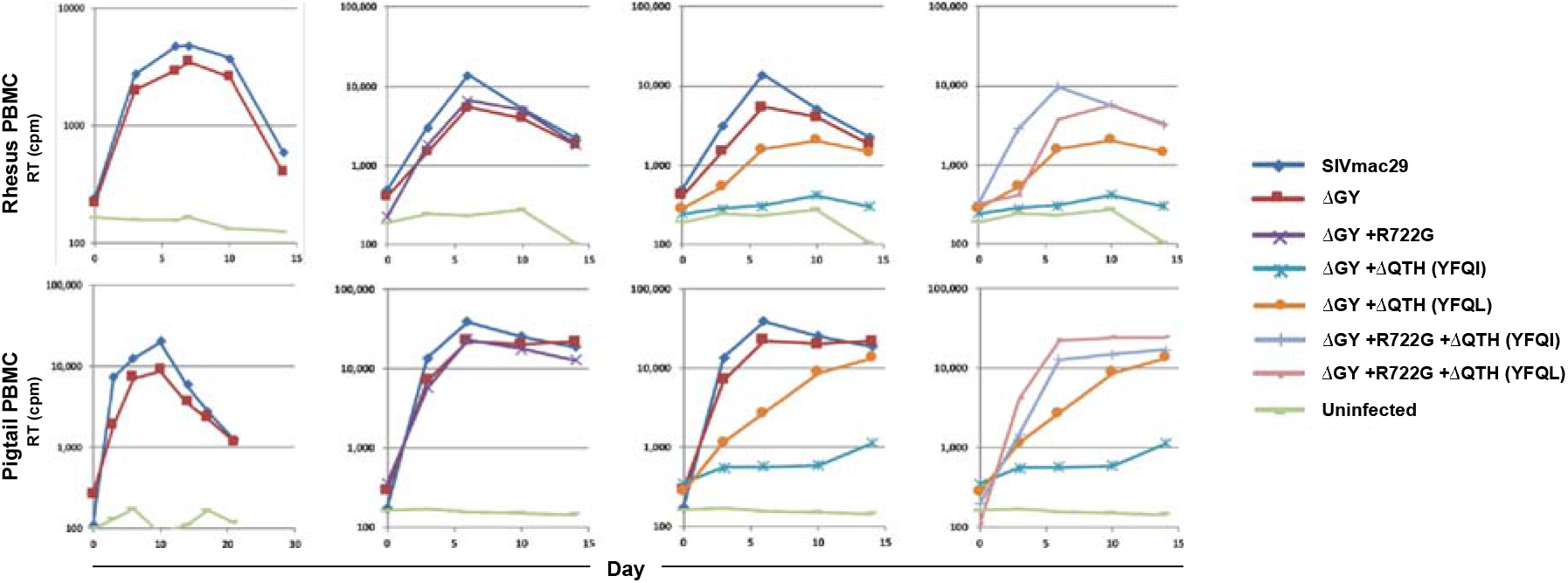
*In vitro* replication of SIVmac239 containing the ΔGY mutation with or without mutations that were acquired *in vivo*. PBMCs from rhesus or pigtail macaques were activated with ConA and IL-2 and infected with SIVmac239, ΔGY, or ΔGY containing the indicated mutations: R722G, ΔQTH creating YFQI (ΔQTH YFQI), ΔQTH creating YFQL (ΔQTH YFQL), or R722G in combination with either ΔQTH (YFQI) or ΔQTH (YFQL). Reverse transcriptase (RT) activity in culture supernatants was measured at the indicated time points. Selected data from 3 separate experiments are shown. SIVmac239ΔGY viruses containing only a ΔQTH mutation replicated poorly but were rescued by addition of R722G.

**Supplemental Figure 5.**
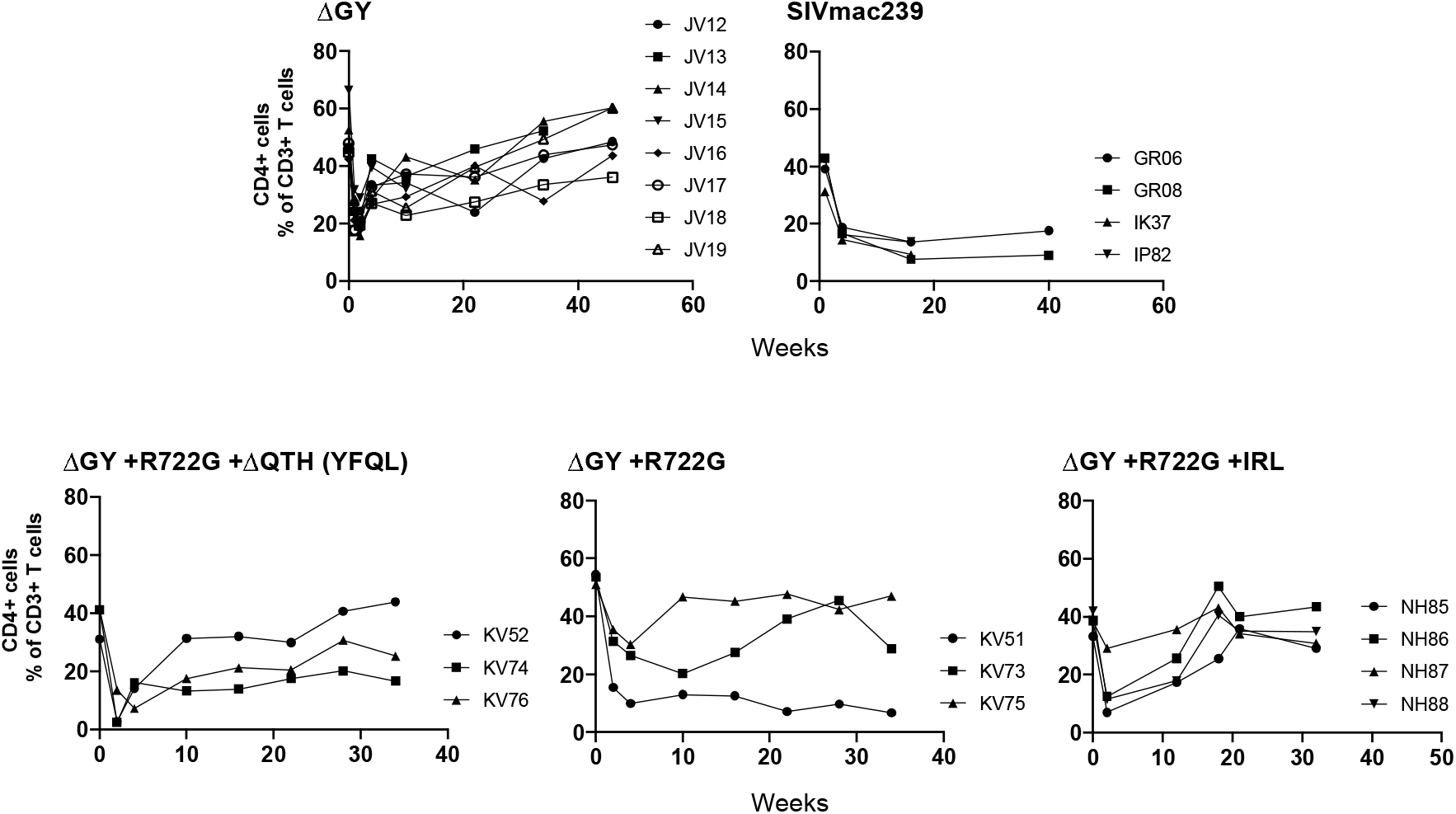
Gut CD4 T cells in pigtail macaques inoculated with SIVmac239ΔGY virus containing R722G with or without a ΔQTH or IRL mutation.

**Supplemental Figure 6.**
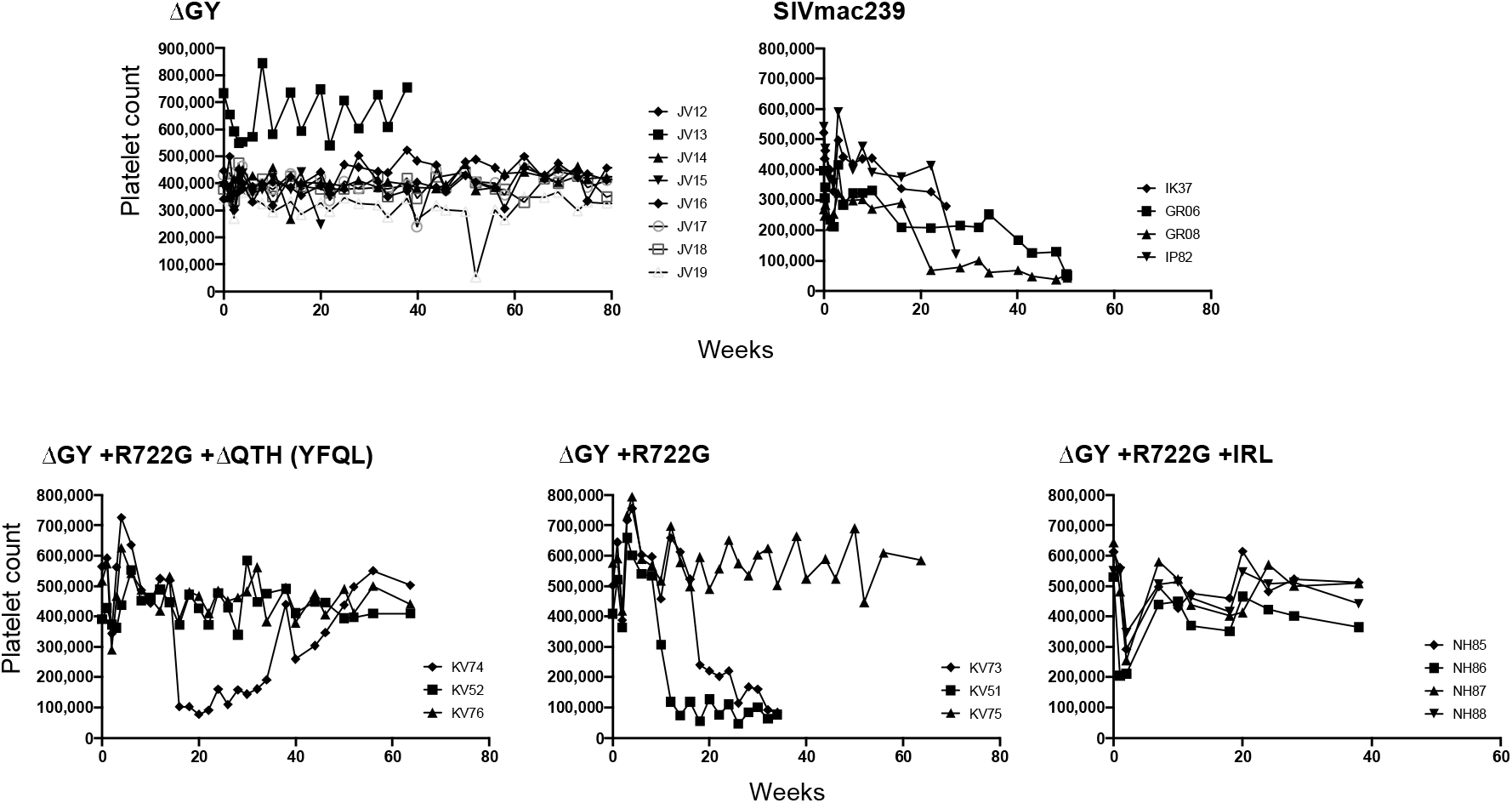
Platelet counts in pigtail macaques inoculated with SIVmac239ΔGY virus containing R722G with or without a ΔQTH or IRL mutation set

**Supplemental Figure 7.**
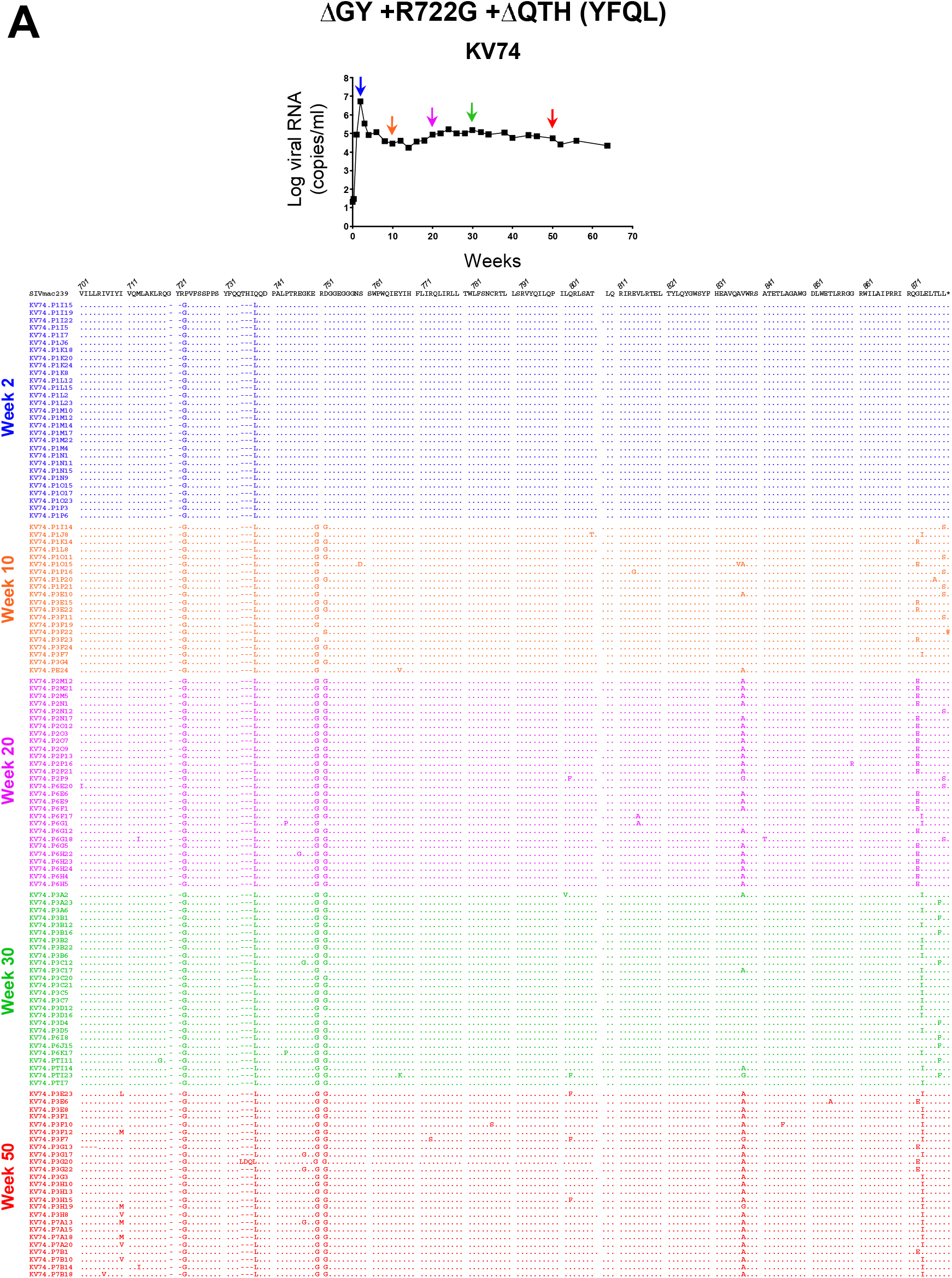
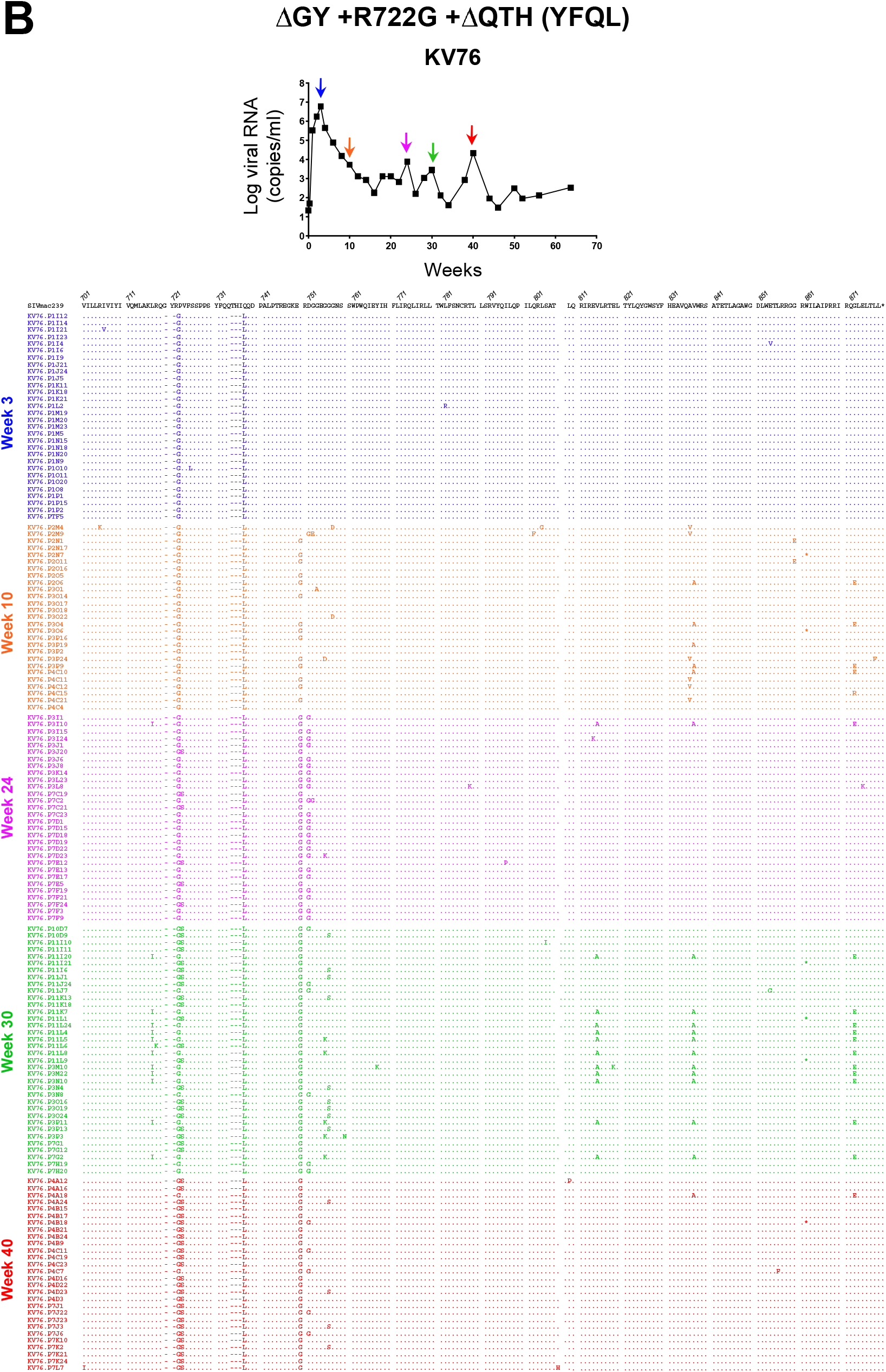
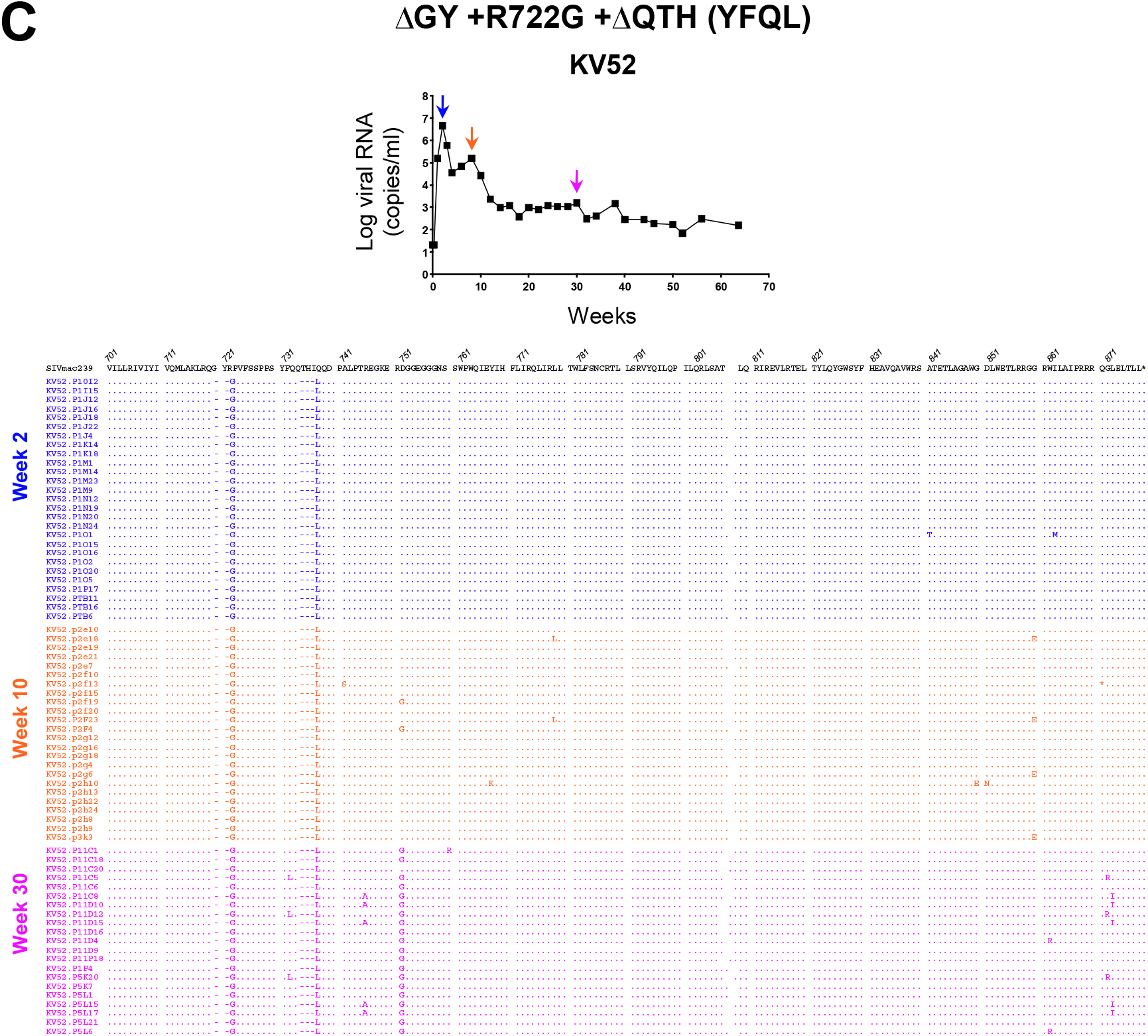
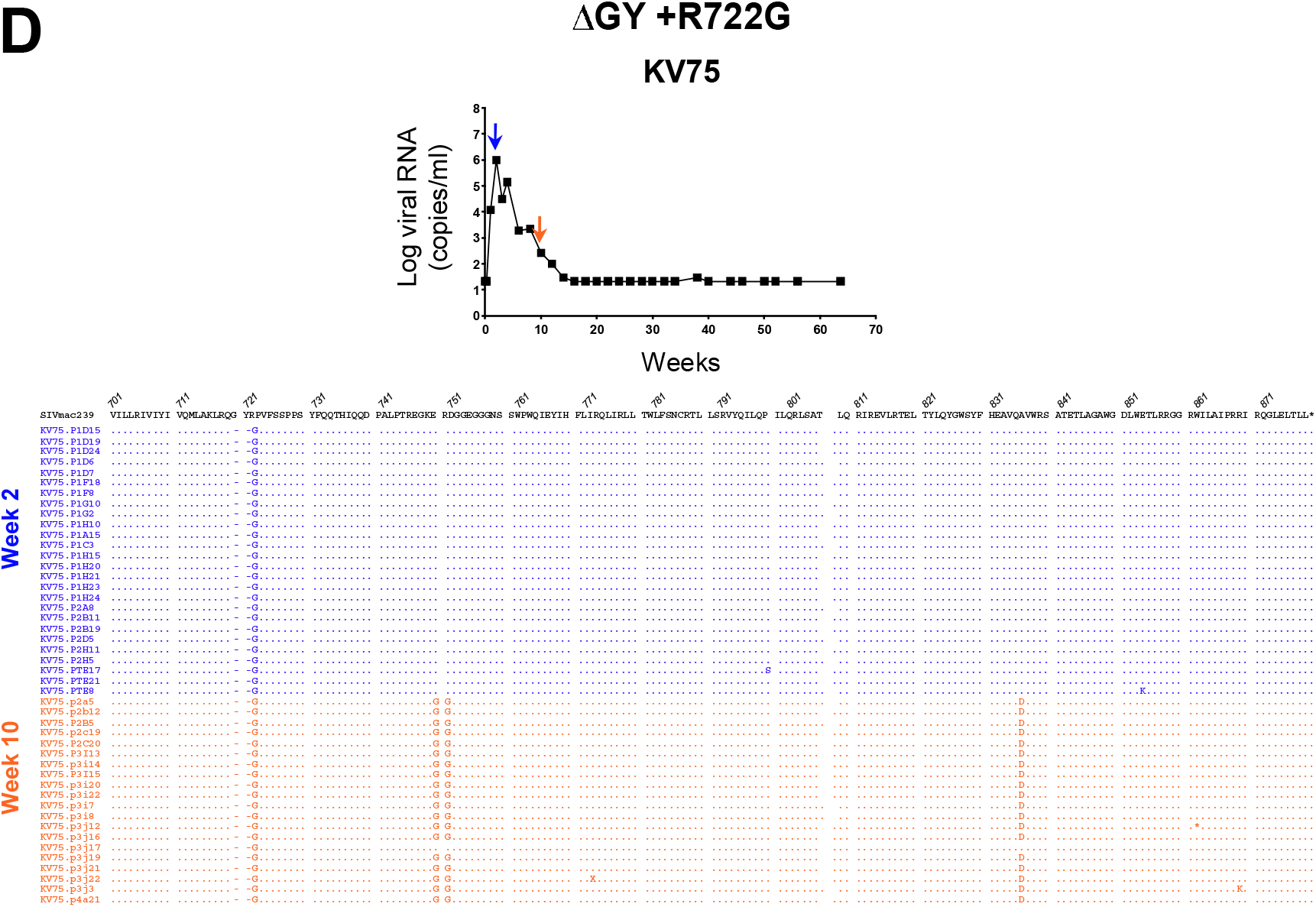
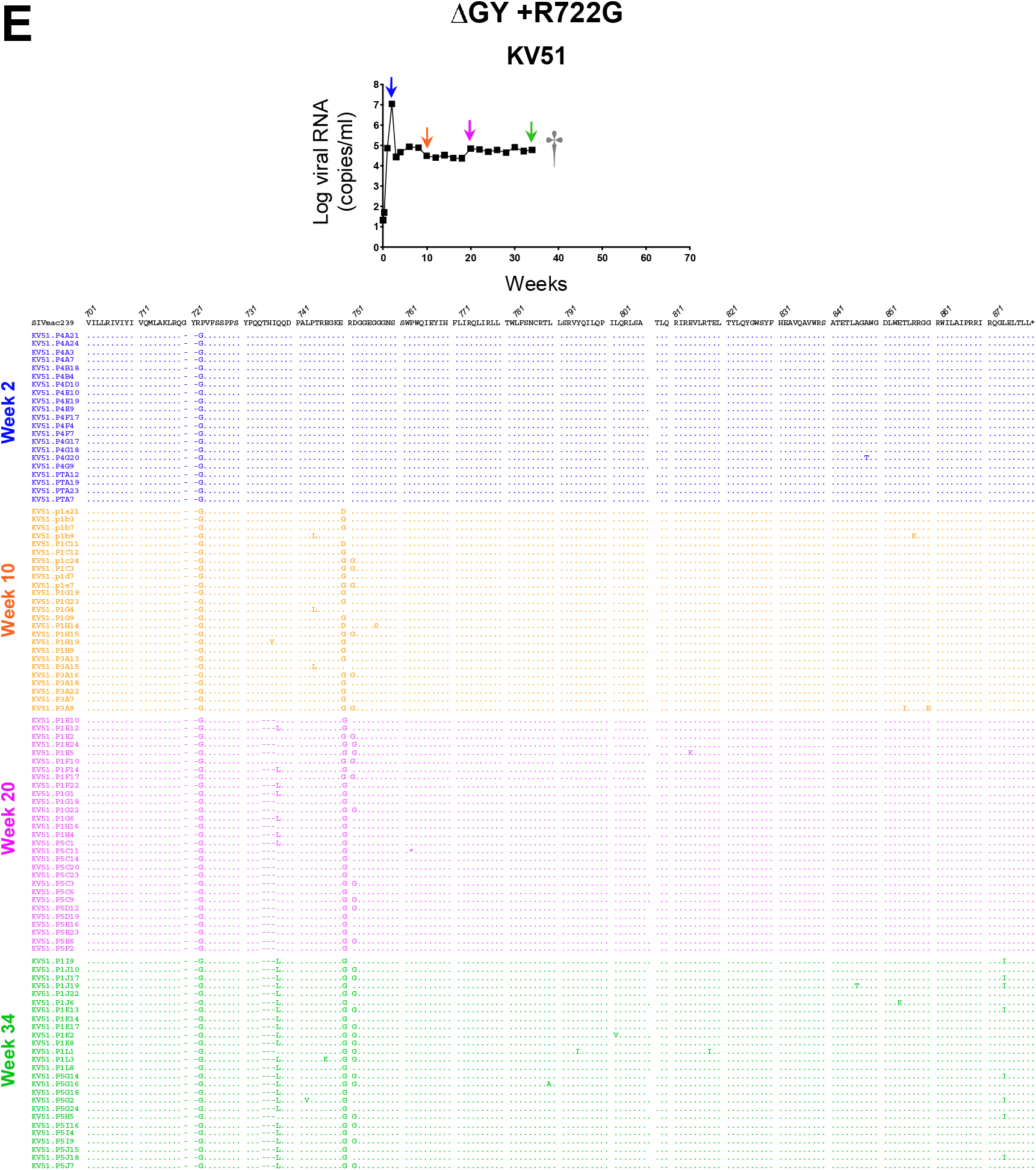
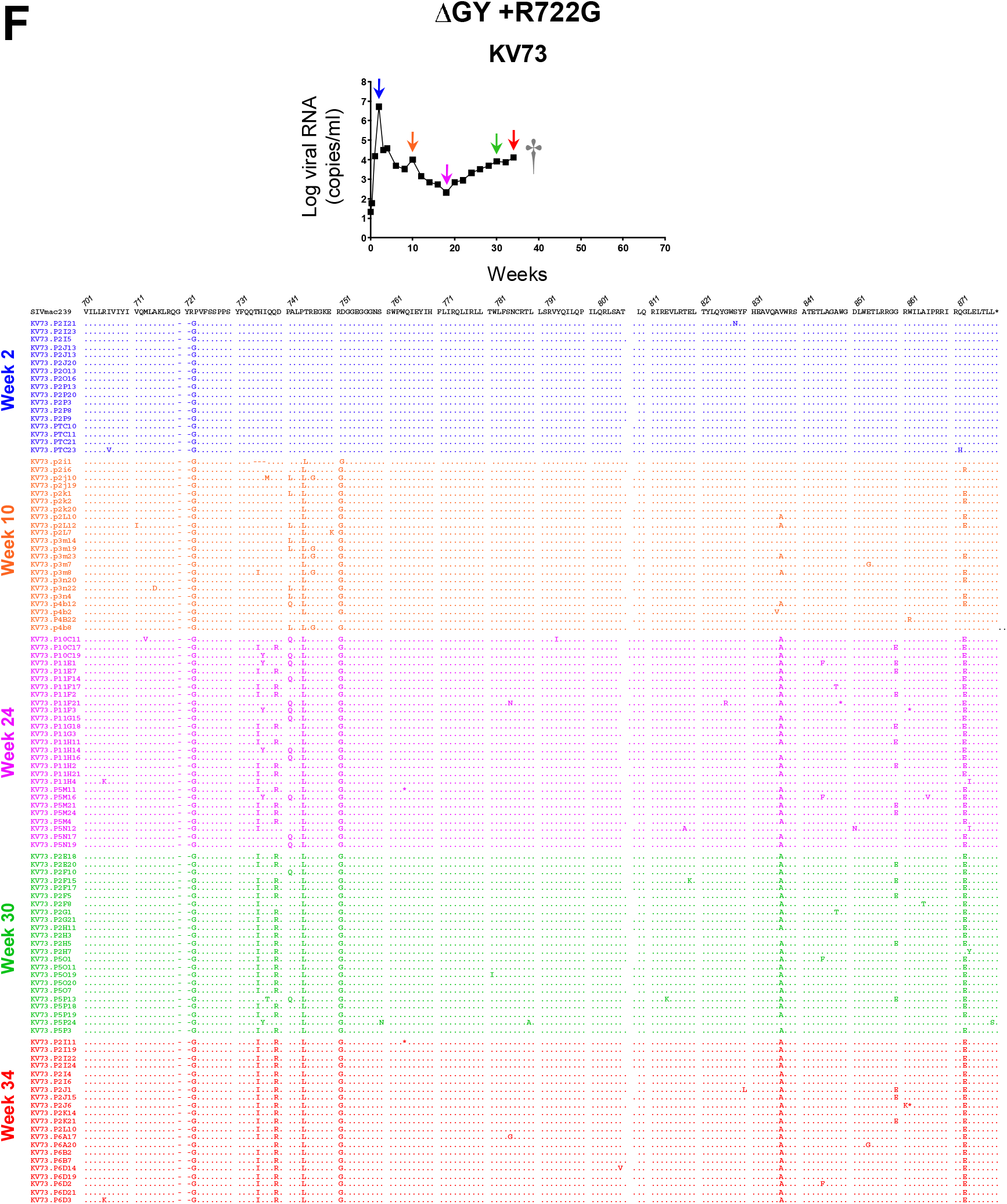
Single genome amplification sequence analyses of plasma viral RNA from pigtail macaques inoculated with SIVmac239ΔGY virus containing R722G with or without a ΔQTH mutation. SGA sequencing of plasma virus from the indicated time points is shown for animals inoculated with SIVmac239ΔGY containing both R722G and ΔQTH mutations (Panels **A**, **B** and **C**), or ΔGY containing R722G (Panels **D**, **E** and **F**). Amino acid sequences (a.a. 701-879) are shown for the Env distal membrane spanning domain and the entire cytoplasmic tail. Amplicons are shown relative to parental SIVmac239 with a.a. identity indicated by a period and deletion mutations indicated by a dash. Plasma viral loads over time for each animal are shown.

**Supplemental Figure 8.**
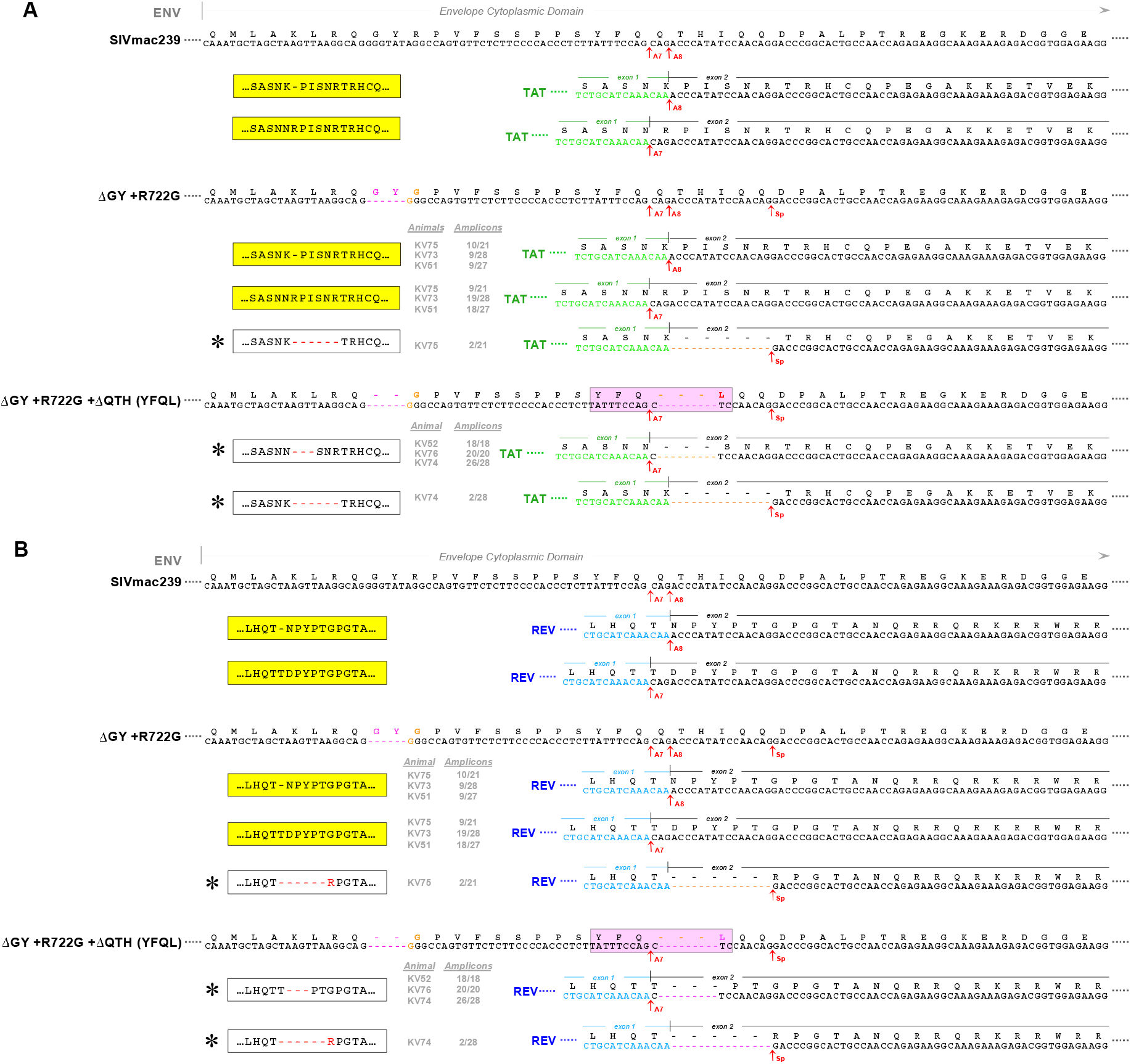
Splicing sites for Rev and Tat mRNAs used *in vivo* during infection with SIVmac239ΔGY variants. mRNA splicing sites for Rev and Tat mRNAs were determined on PBMCs at Day 14 after pigtail macaques were infected with SIVmac239ΔGY containing R722G with or without the ΔQTH mutation that generated the YFQL sequence in Env (**Pink Box**). SGA was used to amplify regions of Rev and Tat mRNAs flanking the predicted splice sites. **(A) Top Panel** shows a.a. and nt sequences for SIVmac239 Env and Tat proteins with *tat* splice acceptor sites A7 and A8 indicated, along with corresponding partial a.a. and nt sequences. Sequences from *tat* exon 1 are shown in **Green**. Amino acid sequences for tat splicing variants are shown (**Yellow Box**). **Middle Panel** shows results for the SIVmac239ΔGY+R722G virus; **Lower Panel** shows results for the SIVmac239ΔGY+R722G+ΔQTH (YFQL) virus. Animal identifiers (see Table 1 and 2) and the number of amplicons exhibiting the indicated splicing pattern relative to the total number of amplicons are shown, as are the corresponding Tat a.a. sequences flanking the splicing sites. Novel variants that were generated are shown and indicated by an asterisk (**\***) **(B)** A similar representation is shown for Rev mRNA splicing patterns for these viruses. Amino acids from Rev exon 1 are shown in **Blue**.

**Supplemental Figure 9.**
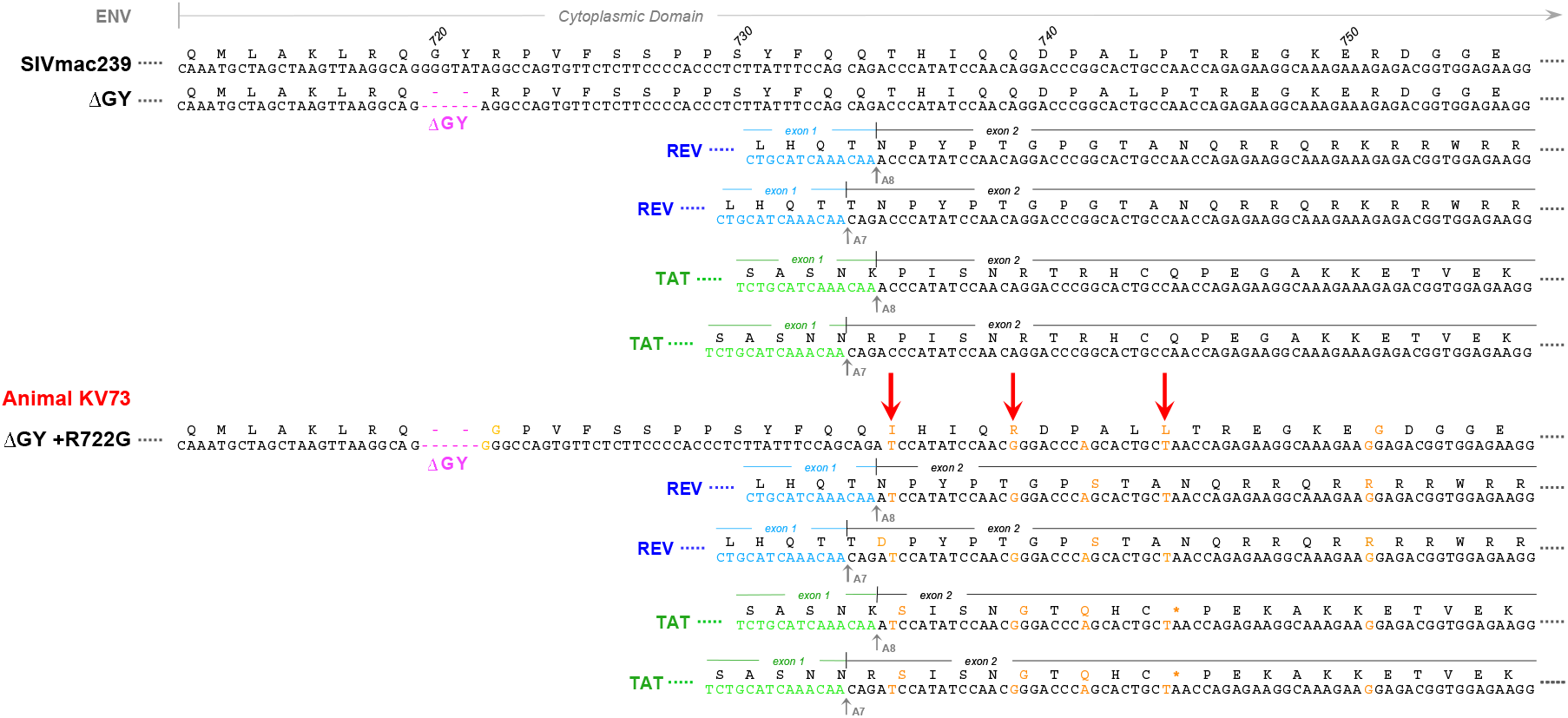
Effects of the IRL Env mutations on *rev* and *tat* open reading frames. **Top Panel** shows a.a. and nt sequences for SIVmac239 and ΔGY Env, Tat, and Rev as shown in Supplemental Figure 1, with known splice acceptor sites indicated and partial a.a. and nt sequences for the 1^st^ exons Rev (**Blue**) and Tat (**Green**). **Bottom Panel** shows point mutations acquired in animal KV73 that was inoculated with SIVmac239ΔGY +R722G (**Magenta**) and that progressed to AIDS (Figure 9 and Supplemental Figure 5C). Nt mutations are indicated in **orange** as are resulting a.a. changes in Env, Rev and Tat. **Red Arrows** show ”IRL” a.a. changes T735I, Q739R, and P744L in Env. A G-to-A nt mutation is also shown that was silent in Env but produced G-to-S and R-to-Q mutations in Rev and Tat, respectively. The C-to-T nt change that produced P744L in Env also generated a stop codon (**\***) in the Tat second exon. The presence of this stop codon was confirmed on mRNAs from macaque PBMCs infected with an SIV that contained the mutations shown above (Brandon Keele, unpublished). The Env R751G fitness mutation is also shown along with its predicted mutation in Rev (43).

**Supplemental Figure 10.**
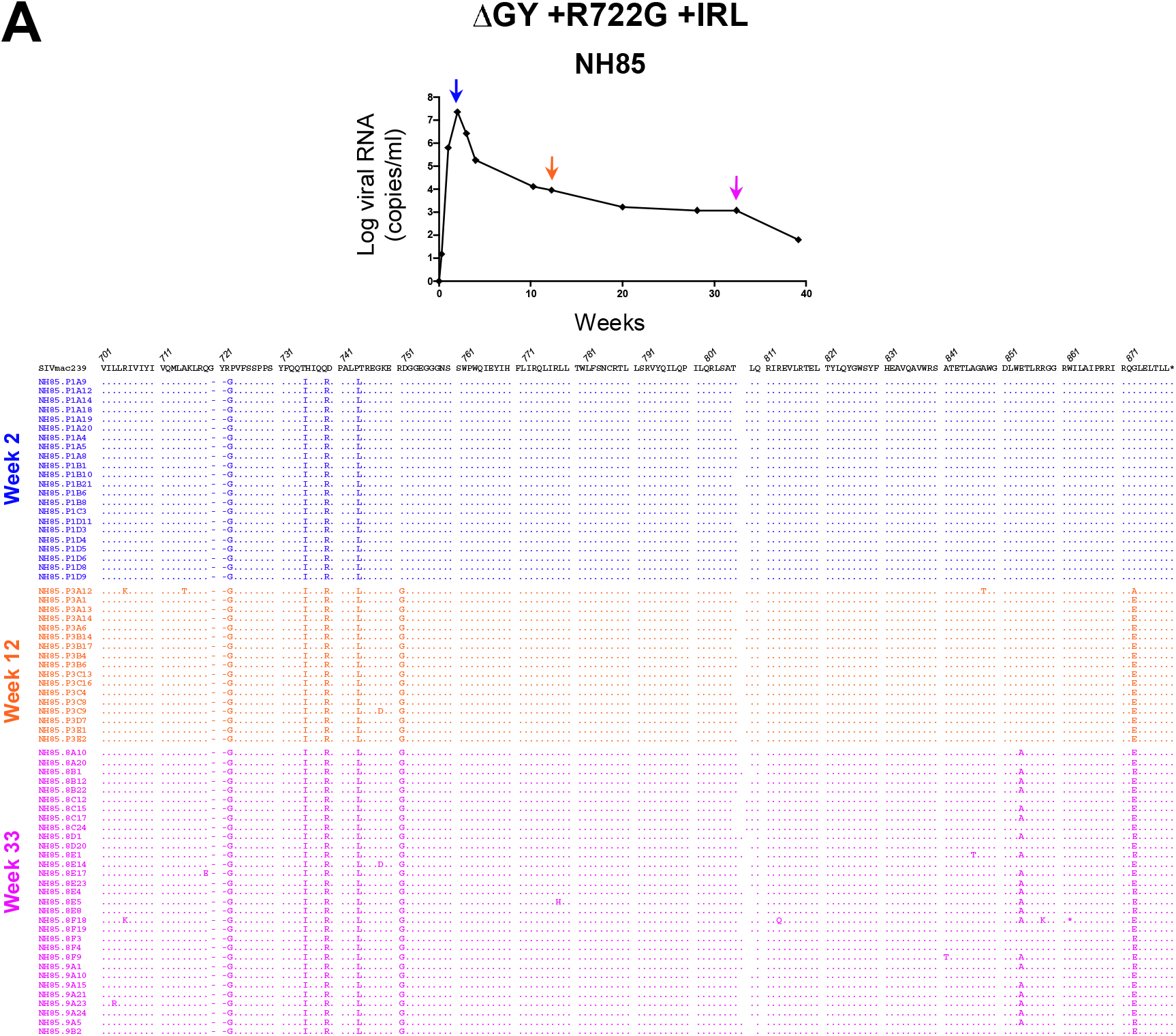
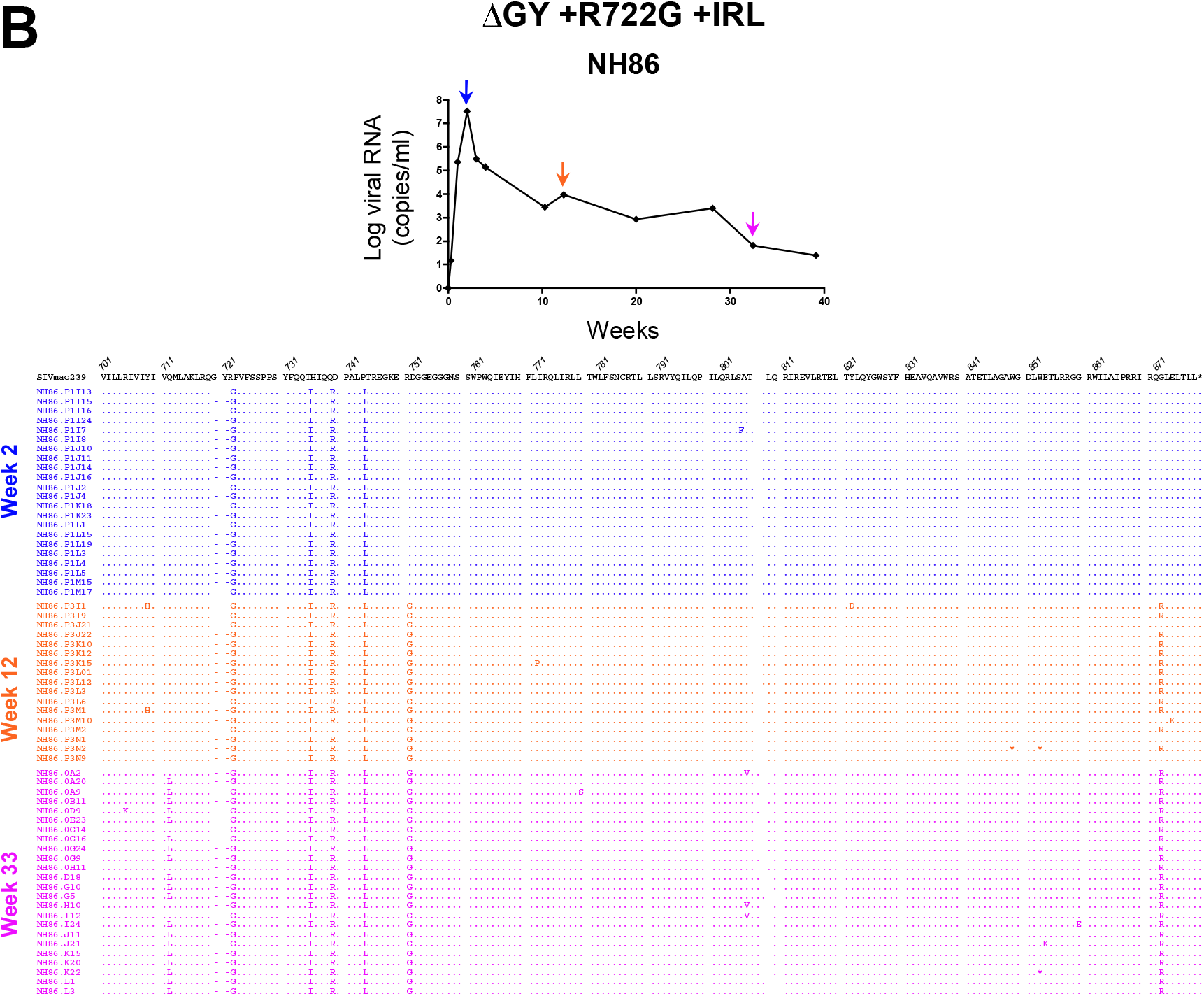
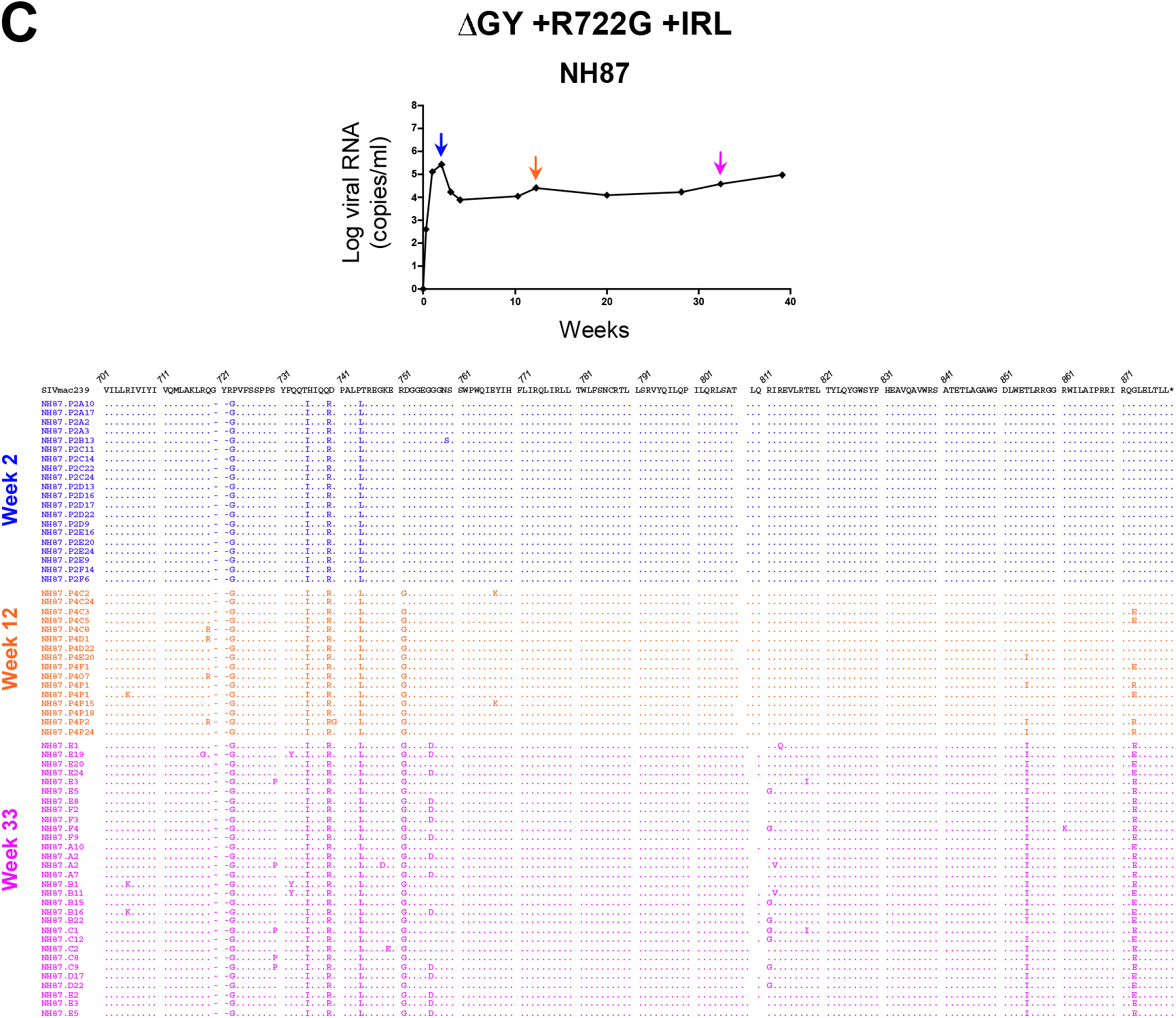
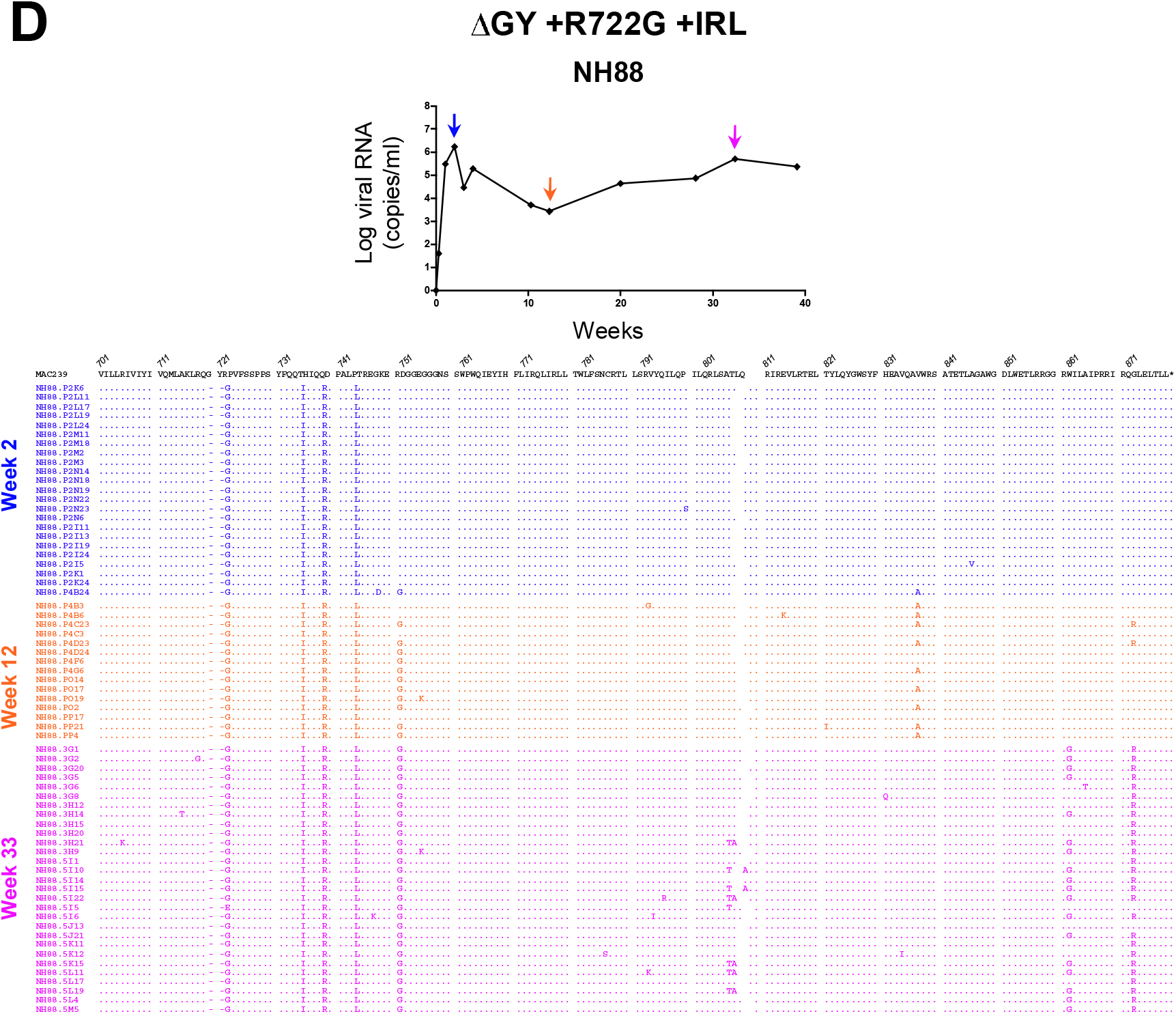
Single genome amplification and sequencing of plasma viral RNA from pigtail macaques inoculated with SIVmac239ΔGY virus containing R722G and the IRL mutation set. SGA of plasma virus from the indicated time points is shown for 4 animals inoculated with SIVmac239ΔGY containing R722G and 3 point mutations (T735I, Q739R, and P744L) shown to confer a new basolateral sorting signal (Figure 7). Amino acid sequences are shown for the Env distal membrane spanning domain and the entire cytoplasmic tail. Amplicons are shown relative to parental SIVmac239 with a.a. identity indicated by a period and deletions indicated by a dash. Plasma viral loads over time for each animal are shown.

## Supplemental Tables

**Supplemental Table 1.**
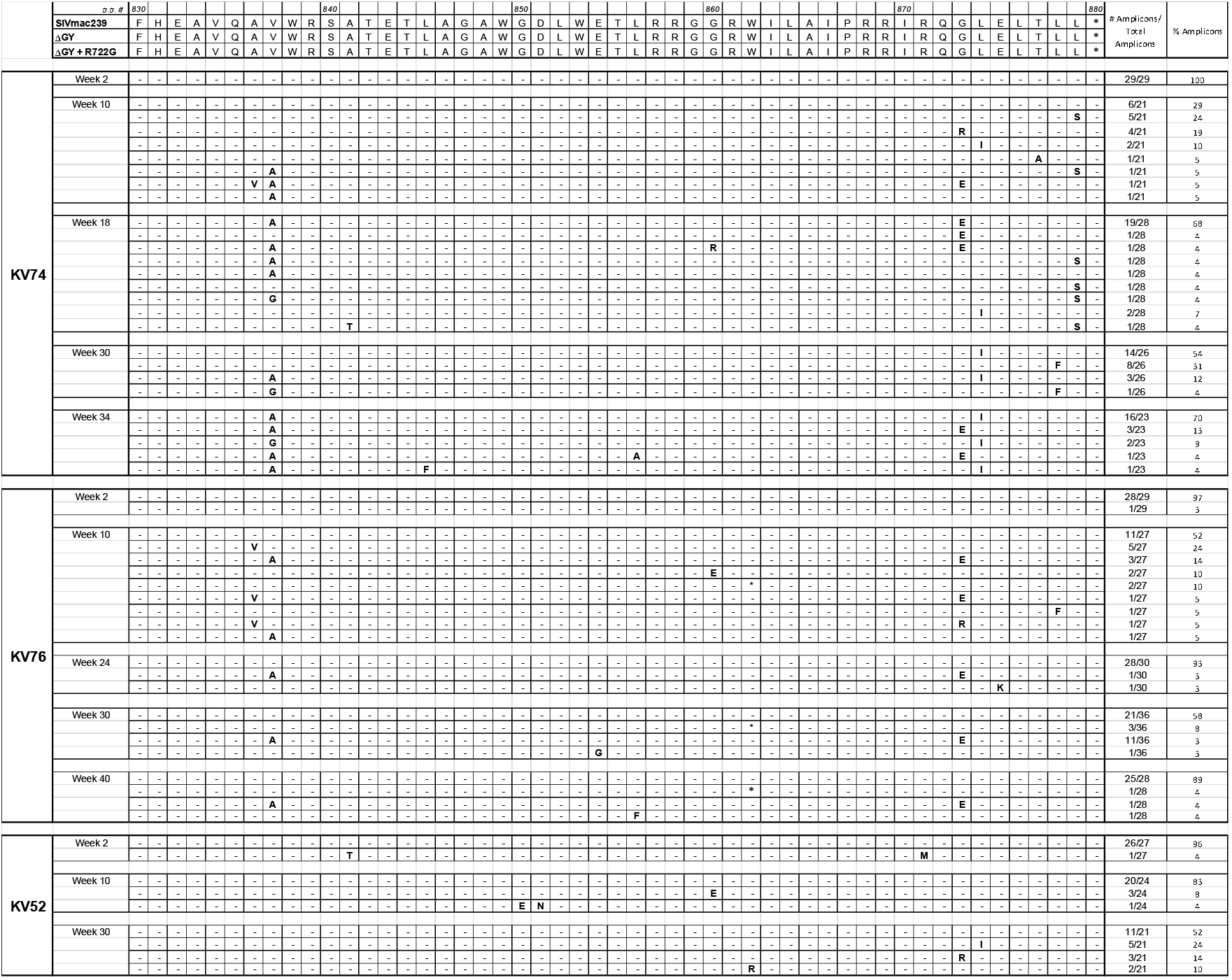
Viral evolution at the Env C-terminus in pigtail macaques inoculated with SIVmac239ΔGY containing the R722G and ΔQTH mutations. Summary of single genome amplification of plasma virus is shown for 3 pigtail macaques inoculated i.v. with the SIVmac239ΔGY+R722G+ΔQTH virus. Sequences for SIVmac239, ΔGY and parental ΔGY +R722G +ΔQTH virus are shown at the top. Results are shown for mutations within a.a. 830 to the C-terminus at position 880. The numbers of amplicons bearing the indicated mutations and the % of amplicons are shown. Dashes indicates identity with ΔGY Env. See Supplemental Figure 7 for a complete listing of sequences of individual amplicons.

**Supplemental Table 2.**
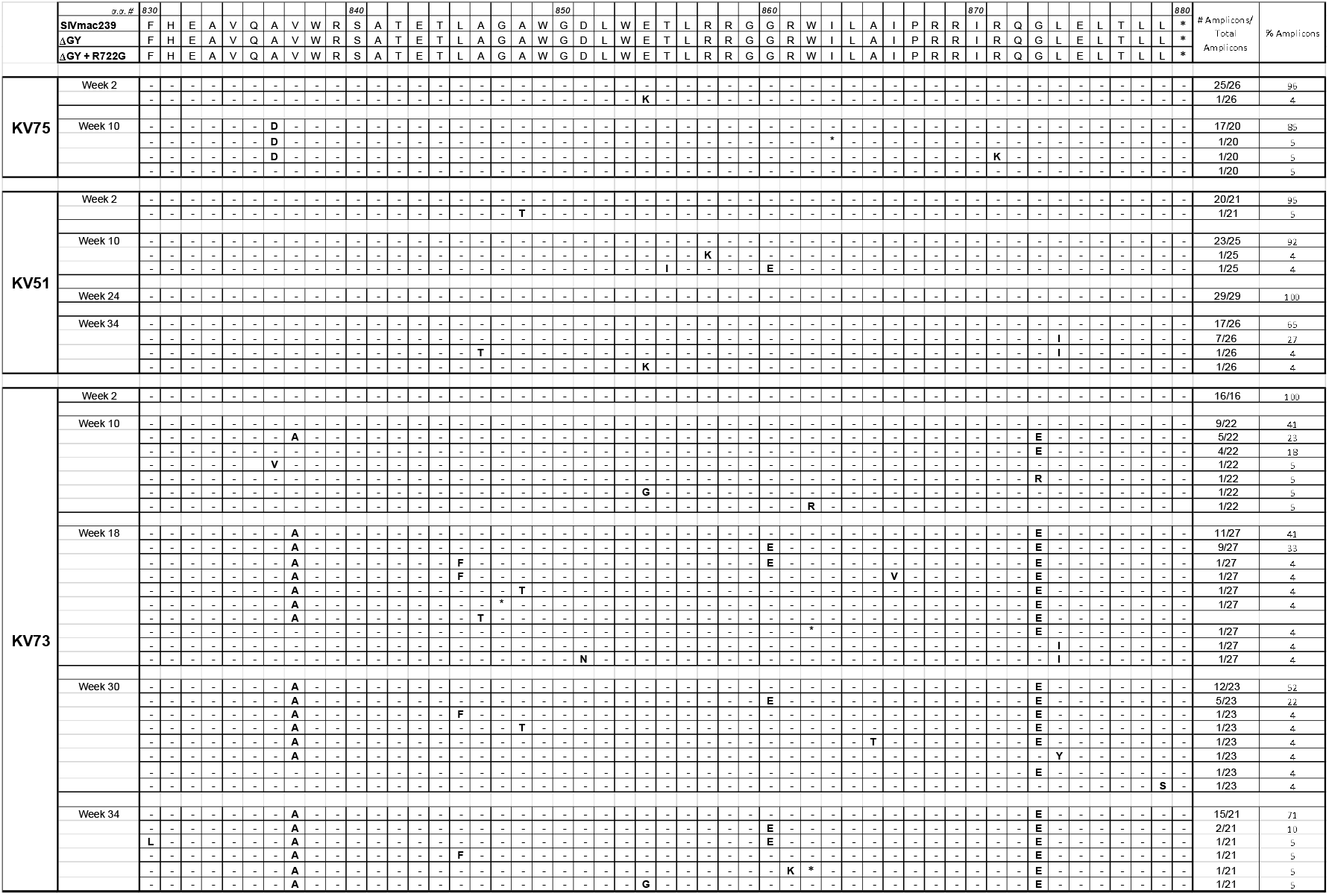
Viral evolution at the Env C-terminus in pigtail macaques inoculated with SIVmac239ΔGY containing the R722G mutation. Summary of single genome amplification analysis of plasma virus performed at the indicated time points for 3 pigtail macaques inoculated i.v. with SIVmac239ΔGY+R722G virus. Results are shown for mutations within a.a. 830 to the C-terminus at position 880 as in Supplemental Table 1. Sequences for parental SIVmac239, ΔGY and ΔGY +R722G Envs are shown at the top. Complete listings of a.a. sequences in the cytoplasmic tail for individual amplicons are shown in Supplemental Figure 7.

**Supplemental Table 3.**
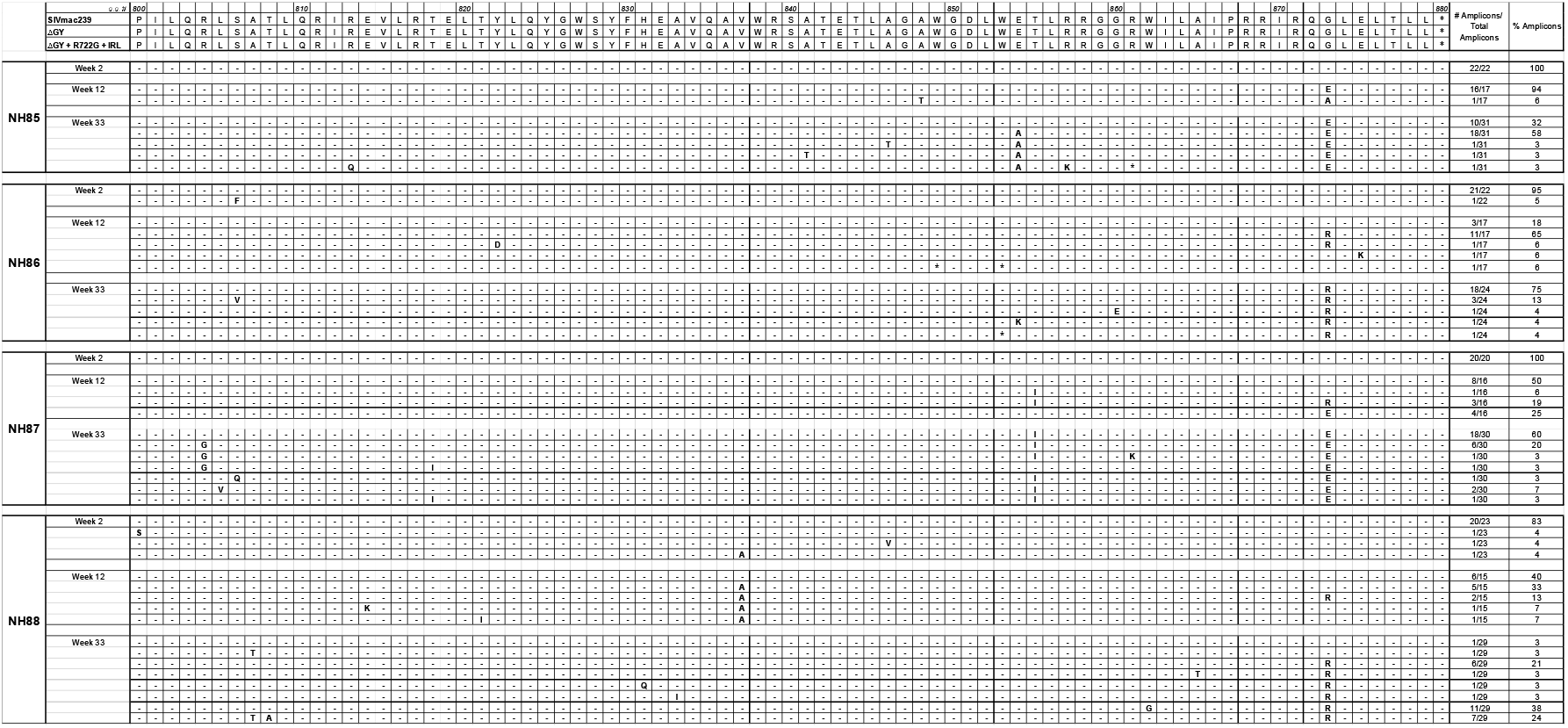
Viral evolution at the Env C-terminus in pigtail macaques inoculated with SIVmac239ΔGY +R722G containing the IRL mutation set. Summary of single genome amplification analysis of plasma virus performed at the indicated time points for 4 pigtail macaques inoculated i.v. with the SIVmac239ΔGY +R722G virus containing the T735I, Q739R and P744L mutations that arose in animal KV53 and conferred a new basolateral sorting signal (see Figure 7). Results are shown for a.a. 800 to the C-terminus at position 880 as in Tables 1 and 2. Sequences for SIVmac239, ΔGY and parental ΔGY +R722G +IRL Envs are shown at the top. Complete listings of a.a. sequences in the cytoplasmic domains for individual amplicons are shown in Supplemental Figure 10.

**Supplemental Table 4.** MHC-I haplotypes of pigtails used in this study. MHC for SIVmac239 and SIVmac239ΔGY infected animals were shown in a previous manuscript (Breed 2015) (n/d, not determined).

| Group | Animal | Sex | Mane-A | Mane-B |
| --- | --- | --- | --- | --- |
| ΔGY +R722G +ΔQTH (YFQL) | KV74 | M | A084b, A093 | B028, B104 |
|  | KV52 | M | n/d | n/d |
|  | KV76 | M | A093, A084b | B028, B028 |
| ΔGY +R722G | KV51 | M | n/d | n/d |
|  | KV73 | M | n/d | n/d |
|  | KV75 | M | A082, A084a | B015b, B028 |
| ΔGY +R722G +IRL | NH85 | M | A010, A084a | B017e, B121 |
|  | NH86 | M | A084a, A038 | B118a, B104b |
|  | NH87 | M | A052a, A084a | B099, B016c |
|  | NH88 | M | A006, A084a | B028, B043 |

